# NOX transmembrane electron transfer is governed by a subtly balanced, self-adjusting charge distribution

**DOI:** 10.1101/2025.03.18.643896

**Authors:** Baptiste Etcheverry, Marc Baaden, Aurélien de la Lande, Fabien Cailliez

**Affiliations:** Institut de Chimie Physique, Université Paris Saclay, CNRS (UMR 8000), 15 avenue Jean Perrin 91405, Orsay, France; Université Paris Cité, CNRS, Laboratoire de Biochimie Théorique, 13 rue Pierre et Marie Curie, 75005, Paris, France

**Author notes:** To whom correspondence should be addressed Corresponding Authors Fabien Cailliez Aurélien de la Lande Marc Baaden.

**Keywords:** Electron transfer, NOX, Molecular dynamics, membrane proteins

## Abstract

NADPH oxidases (NOX) form a family of transmembrane enzymes that catalyze the formation of reactive oxygen species. These are produced thanks to a chain of electron transfers (ET), shuttling electrons from one side of the membrane to the other, using one flavin and two heme cofactors as redox mediators. In this work we investigate the thermodynamics of the electron transfer (ET) between the two hemes contained in the transmembrane domain by means of extensive molecular dynamics simulations. We compare two proteins of the NOX5 isoform, from *homo sapiens* (hNOX5) and from *cylindrospermum stagnale* (csNOX5), a cyanobacteria. We study in detail the influence of both the density of negatively charged lipids in the membrane and of the NOX5 aminoacid sequence on the ET thermodynamic balance. The linear response formalism allows us to decompose the variation in free energy into the individual contributions of the system components (protein, membrane, solvent, etc.). We highlight the major compensatory effects of the various components in the global free energy budget in those complex systems. Although the contributions of the protein or the membrane to the ET thermodynamics can be individually strongly modified by a change in the aminoacid sequence or the membrane composition, they are largely compensated by the rest of the heme environment so that the total free energy is always found to be slightly favorable to the electron transfer. To our knowledge, this study is the first to highlight the effect of membrane charge density on inter-heme ET, providing valuable insights into the molecular mechanisms governing ET catalysis in complex membrane systems.

## INTRODUCTION

Oxidases are enzymes that play a crucial role in living organisms. They are a keystone of ROS (Reactive Oxygen Species) production. ROS are involved in many biological processes such as immune defense in eukaryotes, through the destruction of exogenous pathogens, or cell signaling, notably by ROS concentration in some extracellular environments.^1^ NADPH oxidases (hereafter referred to as NOX) constitute one family of oxidases that have been identified in eukaryote species, dedicated to the production of ROS. Seven isoforms of NOX have been identified (NOX1 to NOX5 and DUOX1/2). Recently, it has been found that NOX-like proteins exist in some prokaryotes.^2,3^ In humans, they are present in several tissues and cell types depending on the isoform: NOX1 is found mostly inside epithelial cells of colon, NOX2 in neutrophils and macrophage cells, NOX3 mainly in the inner ear, while NOX4 has been identified primarily in the renal endothelial cells. Finally, NOX5 and DUOX1/2 are expressed in lymphoid tissues and thyroid cells respectively. The variability in the localization of NOX in eukaryotic organisms is compounded by its uneven distribution within the cell. These transmembrane proteins are present notably through the cytoplasmic membrane, and for some at the level of phagosomal membranes and granules (NOX2). Some NOX family members were also found in the membranes of certain intracellular organelles such as the endoplasmic reticulum (NOX1 and NOX4), mitochondria or even between the inner and outer perinuclear membrane (NOX4).^4^ NOX proteins being widely spread, both at the sub-cellular level and at the scale of an entire organism, their biological impact is large. Production of ROS by NOX is fundamental for the well-functioning of organisms. Its deregulation can induce pathologies such as the chronic granulomatous disease (deficiency in NOX2) or hypertension (NOX1, NOX4 and NOX5).^1^ It is therefore essential to understand the mechanism of ROS production by NOX.

The catalytic core of NOX proteins, depicted in Figure 1, is common to all isoforms. It is composed of a dehydrogenase (DH) domain that binds NADPH substrate and a transmembrane (TM) domain that contains two hemes. An additional flavin cofactor is non-covalently bound to NOX, approximately at the junction of the two domains (in green in Figure 1). ROS production by NOX is achieved through a succession of electron transfer steps, starting with a 2-electron reduction of flavin by NADPH. The electrons are transferred to the closest heme (Heme1, shown in red in Figure 1) and then towards the second (distal) heme (Heme2, in blue in Figure 1). The final oxygen reduction occurs at Heme2 and leads to the formation of superoxide anion (in NOX1, NOX2, NOX3, and NOX5) or hydrogen peroxide (in NOX4 and DUOX1/2). In addition to these successive reactions, NOX function requires a structural activation step (except for NOX4 which is constitutively active). This activation is supposed to trigger a conformational modification to bring together NADPH and flavin, allowing the hydride transfer. The activation mechanism is different among the isoforms. NOX1, NOX2 and NOX3 assume their active form through the assembly of cytosolic proteins around the dehydrogenase domain, while NOX5 and DUOX1/2 activity is regulated by the binding of calcium ions to an EF-hand subdomain.^5^

**Figure 1.**
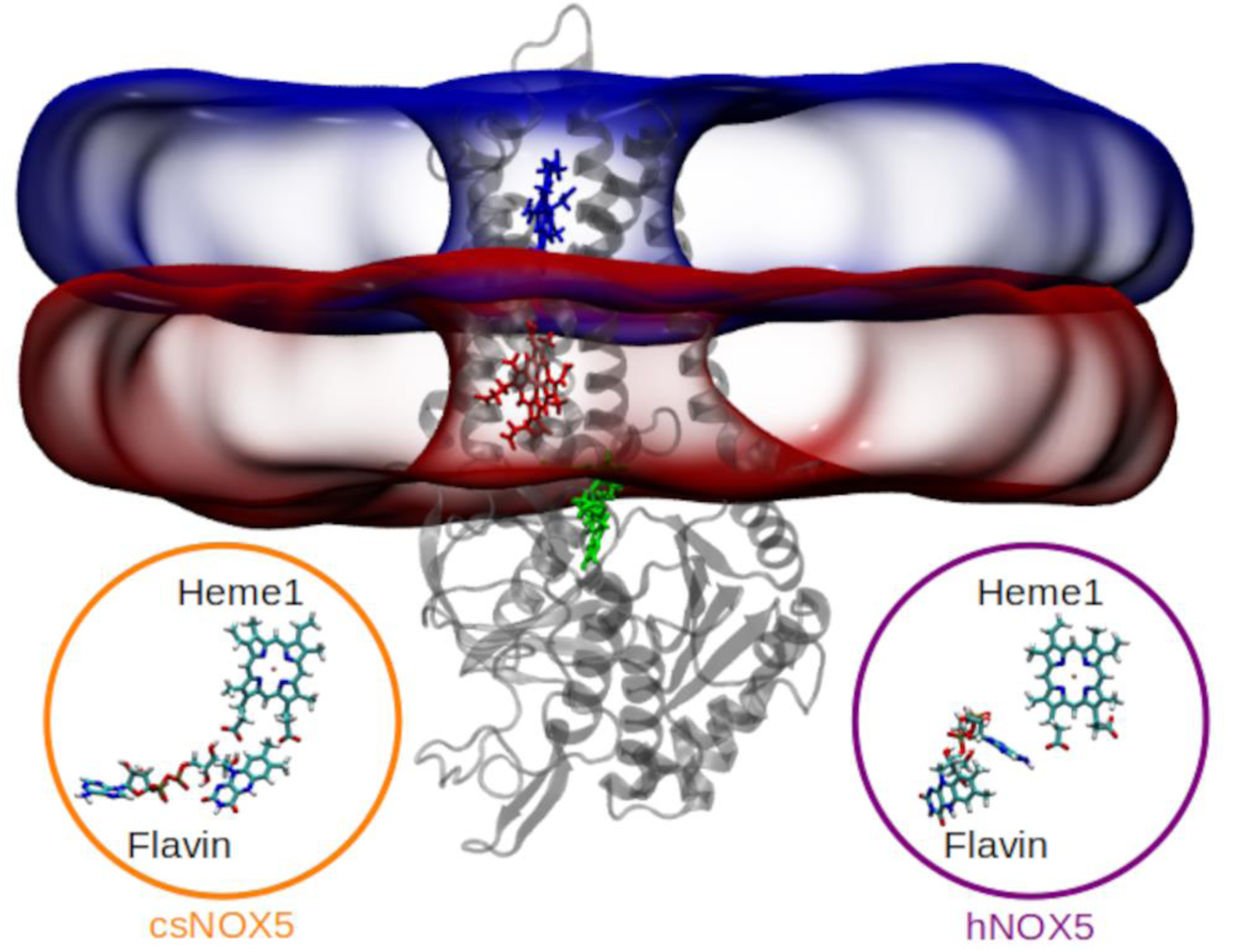
Illustration of the hNOX5 model inserted in a lipidic membrane showing the localization of the three cofactors: flavin (in green), Heme1 (in red), and Heme2 (in blue). The catalytic core is shown in grey using a schematic representation. The surfaces of the 2 leaflets of the membrane are represented with the Quicksurf representation of VMD^6^, in blue for the “upper” leaflet and in red for the “lower” leaflet. At the bottom of the figure, in the circles, a zoom on the relative orientation of the flavin with respect to Heme1 is shown in the csNOX5 model (on the left) and in the hNOX5 model (on the right)

The first experimental model of a full NOX isoform has been determined by X-ray crystallography of the *cylindrospermum stagnale* NOX5 protein (csNOX5).^7^ The DH and TM domains were crystallized separately, and a model of the full structure was proposed by docking of the two domains. This work revealed the position of the two heme cofactors at the heart of the TM domain and proposed a geometric arrangement of the DH and TM domains compatible with a direct electron transfer from the flavin to the first heme. A similar model has been built by our team and its structural stability was tested by molecular dynamics simulations in a previous work^8^. Since then, structures of various other proteins from the NOX family have been solved experimentally, most of them using cryo-electron microscopy (Cryo-EM): NOX2 from human^9–11^, human NOX5^12^, mouse and human DUOX1^13,14^, a NOX homolog from *Streptococcus pneumoniae* (SpNOX)^3,15^. This panel of structures corresponds to various states of activation of NOX. For instance, the Cryo-EM structure of human DUOX1 has been determined in two states^14^, one supposedly active, the other one a rested state of the enzyme. Likewise, structures of the DUOX1 enzyme from mouse were published by Sun in 2021 in their low and high calcium state, showing notably differences in the EF-hand structure and orientation^13^. More recently, NOX2 structures have been determined in the resting and in the activated states^11^. The accumulated structural information makes it possible to elaborate hypotheses about the conformational modification happening during activation^3^. Regarding the electron transfer steps, the available experimental data confirm that the TM domain structure is highly conserved in the family with a similar arrangement of the two hemes. They show a rather similar arrangement of the flavin with respect to the first heme, in which the adenosine moiety of flavin is closer to the heme than the isoalloxazine ring.

Molecular simulation is a powerful way to complement experimental approaches. It allows to explore system dynamics and to compute thermodynamic observables related to electron transfer in various controlled scenarios^16,17^. Our current knowledge of electron transport in multi-heme protein has been largely assessed by advanced computer simulations^18^. Based on the csNOX5 structures of DH and TM domains, we reported the very first study of the inter-heme electron transfer in NOX proteins in 2021^8^. In this work, we built a model of a full NOX protein catalytic core (DH+TM) inserted in a lipidic membrane composed of a mixture of 1-palmitoyl-2-oleoyl-sn-glycero-3-phosphoethanolamine (POPE) and 1-Palmitoyl-2-oleoyl-sn-glycero-3-(phosphor-rac-(1-glycerol)) (POPG) lipids with a POPE:POPG ratio of 1:4. We then computed the free energy (Δ*G*) of the inter-heme electron transfer (ET), obtaining a value of around -0.3 eV for Heme1-to-Heme2 ET. Although consistent with the directionality of the electron transfer in NOX, this favorable Δ*G* value contrasts with the unique experimental measurement of heme redox potentials in NOX2^19,20^ (−225 mV for Heme1 and -265 mV for Heme2 at pH 7) that would indicate unfavorable ET. Actually, our previous computations showed that besides the redox cofactors and the protein itself, the lipid membrane, the water environment and the counterions largely contribute to determine the Δ*G* value, making direct comparison with the available experimental data delicate. Furthermore, some theoretical studies revealed that protein dynamics can be impacted by the membrane composition^21^. Membrane composition can have an influence on protein assembly at the surface of the bilayer^22^. In the same trend, some experimental studies demonstrated the impact of cytoplasmic or phagosomal membrane charges, as well as asymmetric leaflets on enzyme activity, and more precisely on NOX proteins^23–25^. Finally, the composition of the membrane is subject to modifications during NOX2 activation^26,27^. This ensemble of information has motivated us to explore in depth the interplay between the membrane composition, in particular its electric charge state and the energetics of the electron transfer. This is an aspect that to our point of view has not yet been sufficiently addressed in molecular simulations dealing with ET in membrane proteins^8,28–32^.

Here, we go beyond our previous work and bring further characterization of inter-heme ET thermodynamics in NOX systems. Our goal in this work is twofold. First, we re-evaluate the relevance of our first csNOX5 model by comparing it to the more recent experimental structures of human NOX5 (hNOX5)^12^. Second, we aim at understanding if and how the environment modulates the thermodynamics of the Heme1-to-Heme2 ET. We are particularly interested in deciphering the role of the lipid membrane composition.

To this aim, we carried out extensive molecular simulations of NOX5 from two species (csNOX5 and hNOX5) in various conditions. These two models allow us to monitor the influence of the protein sequence on the inter-heme electron transfer. We inserted these NOX models in two lipidic membranes with two very different compositions, varying the POPE:POPG ratio. Doing so, we can evaluate the influence of the membrane on the structure and dynamics of our models, as well as on the thermodynamics of the inter-heme ET. We have used multiple hundreds of nanoseconds-long molecular dynamics simulations of those systems in the two redox states associated with the inter-heme electron transfer to compute ET reaction free energy, using a protocol detailed in the Computational Methods section. In the first part of the Results section, we analyze the structure and dynamics of the NOX models to assess their stability. We then present our calculations of the free energy of the inter-heme electron transfer for all models, with a particular focus on the contribution and detailed balance of the various components of the system. The dynamical and thermodynamical results are then discussed.

## COMPUTATIONAL METHODS

### csNOX5 and hNOX5 models

For csNOX5, we used a model built from the experimental structure of the separate TM (5O0T protein data bank (PDB) code, 2.0 Å resolution) and DH (5O0X PDB code, 2.2 Å resolution) domains of csNOX5^7^. Details of the construction of the model are given in our previous work. As a starting point of the molecular simulations described in this work, we have extracted a representative structure from one of our previous molecular dynamics simulations (centroid of the most populated cluster)^8^.

The hNOX5 model is based on the recent Cryo-EM structure of Cui *et al.*^12^ (PDB code 8U85). This structure was obtained at low calcium concentration and thus corresponds to a non-activated NOX5 system. The experimental structure of NOX5 is dimeric but we chose to keep only one monomer of hNOX5 in our simulations for the sake of comparison with the csNOX5 system. For the same reason, we have removed the EF-hand domain (residues 1 to 179) which is absent in the csNOX5 model. Missing residues in the experimental structure have been rebuilt using the ColabFold software, an alternative version of the Alphafold2 tool^33^. The protocol used was the following: we used ColabFold to predict the structure of hNOX5 using the Cryo-EM structure as template and 18 cycles of refinement. Despite the use of a template, the final predicted structure’s alignment deviated from the experimental structure. However, local alignments on the helices flanking the missing residues were satisfactory. We thus chose to rely on the experimental atomic positions when available and take the missing atomic positions from the ColabFold model, locally aligned to the experimental structure to build a consolidated model. Some short molecular mechanics energy minimizations have been performed to avoid side chain clashes and finally obtain the initial protein structure used for hNOX5 simulations.

csNOX5 and hNOX5 proteins, originating from different organisms, do not share the same primary sequence (Figure S1). They share a total sequence identity of 38%. The major differences arise in the EF-hand domain and the TM domain (34% of identity). However, a lot of amino-acid patterns remain preserved around the hemes and flavin cofactors. Despite this low sequence identity, the structure of the DH and TM domains are close. We notice two major structural differences. The first one is the DH domain contraction. The DH domain is composed of a flavin binding domain (FBD) and a NADPH binding domain (NBD). According to Liu *et al.*^11^, the relative position of these two domains is a marker of the state of activation of NOX2. They use the distance between the Cα atoms of Arg356 (in the FBD) and Cys537 (in the NBD) residues to measure the contraction of the DH domain. Following their proposition, we use the same distance between the corresponding conserved residues in csNOX5 (Arg270 and Cys444) and hNOX5 (Arg509 and Cys740) to describe the contraction of the DH domain (see Figure 2a). This measure is 13.1 Å and 18.7 Å in the initial models of respectively csNOX5 and hNOX5, prior to molecular dynamics simulation. The contraction of csNOX5 is much higher than the one of hNOX5.

**Figure 2.**
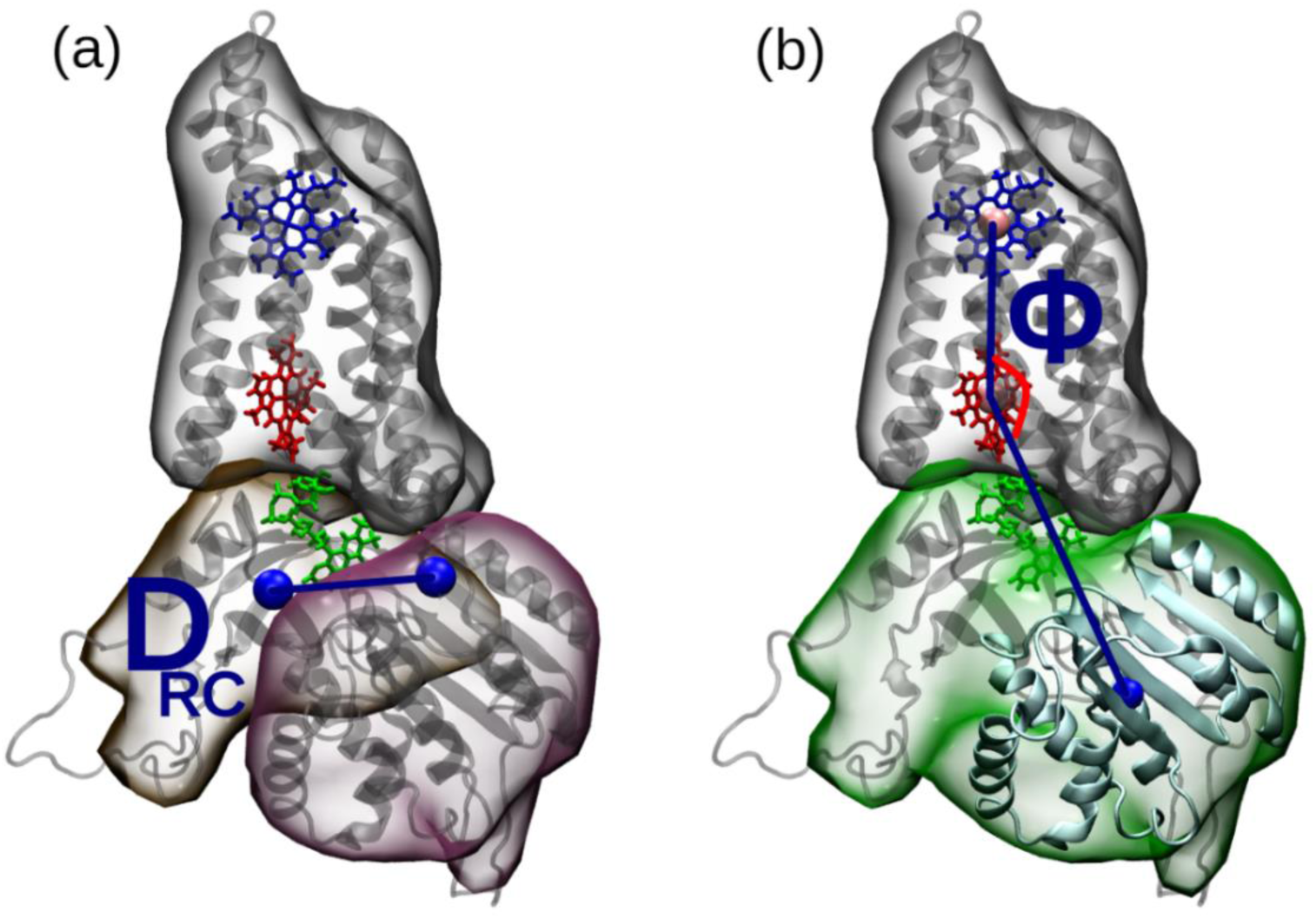
Geometric features characterizing NOX5 conformations. (a) the distance between Cα atoms of residues Arg509 and Cys740 (in csNOX5 numbering), highlighted by blue spheres defines the contraction of the DH domain. The TM, FBD and NBD domains are depicted with grey, yellow, and pink transparent surfaces respectively; (b) ϕ is a hinge angle between the TM and the DH domain. The two heme irons are shown with pink spheres. The TM, and DH domains are depicted with respectively a grey and green transparent surface.

The second major difference between our two models lies in the orientation of the flavin cofactor with respect to Heme1. This difference of orientation in the two models is illustrated in Figure 1. In the csNOX5 model, the isoalloxazine ring of the flavin faces Heme1. Note that this orientation originates from restraints imposed in the process of construction of the model, described in Wu *et al.*^8^, but has been conserved during the 300ns-long MD simulations in our previous work. In that orientation the isoalloxazine ring is stacked with the C-terminal residue of the DH domain (Phe469). In the hNOX5 structure, it is the adenosine moiety of the flavin that is closer to Heme1. There is no π-stacking interaction with Phe765, the C-terminal residue of the DH domain, but the isoalloxazine ring is hydrogen-bonded with 4 residues in its surrounding (Arg509, His507, Thr594 and Thr493).

### Preparation of the full system

The protein’s amino-acids residues have been assigned a standard protonation state at pH 7 with all histidine residues singly protonated at the Nd site. We have used CHARMM-GUI^34–36^ to create the lipidic membranes around the NOX5 models. Two different membrane compositions have been set up: one highly anionic membrane with a 4:1 POPG:POPE ratio (hereafter referred to as mbH membrane, H standing for “Highly”), and a more weakly charged one with a roughly 1:4 POPG:POPE ratio (referred to as mbW membrane, W standing for “Weakly”). Prior to insertion in a lipid membrane, the protein transmembrane domain was oriented using the OPM (orientations of proteins in membranes) web server^37^. The protein and membrane systems were then solvated using TIP3P^38^ water molecules in a hexagonal prism unit cell. Finally, sodium and chloride ions were added to neutralize the systems and achieve an ionic concentration of about 0.15M. Details of the initial composition of each system are given in Table 1.

**Table 1.** Details of the simulations’ setup. For each simulated system are given the absolute numbers of POPE and POPG lipids in each leaflet (Low and Up), the POPE:POPG ratio for the entire membrane and within each leaflet, the number of sodium and chloride counterions, and the charge states of the two hemes, as well as an acronym for reference.

| NOX5 model | Lipid leaflets composition | POPE:POPG ratio | Counterions | Redox state | System name |
| --- | --- | --- | --- | --- | --- |
| csNO5 | Low:<br>POPE 164 / POPG 31<br>Up:<br>POPE 172 / POPG 38 | Total: 4.9:1<br>Low: 5.2:1<br>Up: 4.5:1 | 164 Na<sup>+</sup><br>102 Cl<sup>-</sup> | Heme1-Fe<sup>2+</sup><br>Heme2-Fe<sup>3+</sup> | cs_mbW_st1 |
|  |  |  |  | Heme1-Fe<sup>3+</sup><br>Heme2-Fe<sup>2+</sup> | cs_mbW_st2 |
|  | Low:<br>POPE 40 / POPG 155<br>Up:<br>POPE 42 / POPG 168 | Total: 1:4<br>Low: 1:4<br>Up: 1:4 | 417 Na<sup>+</sup><br>101 Cl<sup>-</sup> | Heme1-Fe<sup>2+</sup><br>Heme2-Fe<sup>3+</sup> | cs_mbH_st1 |
|  |  |  |  | Heme1-Fe<sup>3+</sup><br>Heme2-Fe<sup>2+</sup> | cs_mbH_st2 |
| hNOX5 | Low:<br>POPE 160 / POPG 40<br>Up:<br>POPE 172 / POPG 38 | Total: 4.3:1<br>Low: 4:1<br>Up: 4.5:1 | 175 Na<sup>+</sup><br>117 Cl<sup>-</sup> | Heme1-Fe<sup>2+</sup><br>Heme2-Fe<sup>3+</sup> | h_mbW_st1 |
|  |  |  |  | Heme1-Fe<sup>3+</sup><br>Heme2-Fe<sup>2+</sup> | h_mbW_st2 |
|  | Low:<br>POPE 40 / POPG 160<br>Up:<br>POPE 43 / POPG 167 | Total: 1:4<br>Low: 1:4<br>Up: 1:4 | 424 Na<sup>+</sup><br>117 Cl<sup>-</sup> | Heme1-Fe<sup>2+</sup><br>Heme2-Fe<sup>3+</sup> | h_mbH_st1 |
|  |  |  |  | Heme1-Fe<sup>3+</sup><br>Heme2-Fe<sup>2+</sup> | h_mbH_st2 |

### Molecular dynamics simulations

The CHARMM36 forcefield for proteins, lipids and counterions^36^ has been used in combination with the TIP3P forcefield for water^38^. The parameters for the two hemes in the two redox states (ferric and ferrous) were taken from our previous work^8^. The flavin cofactor has been modeled in its semi-reduced FADH° state using the recent parameters of Aleksandrov, compatible with the CHARMM forcefield^39^.

For the eight systems defined in Table 1 we used the standard protocol of equilibration provided by CHARMM-GUI. It consists of 6 successive steps of 25, 25, 25, 200, 200, and 200 ps of molecular dynamics simulations in the isothermal-isobaric ensemble (NPT) using periodic boundary conditions at 303K and 1 bar. During this procedure, harmonic restraints were put on protein backbone atoms, lipids’ centers of mass and heme and flavin cofactors and gradually decreased. Harmonic forces were set to 10.0, 5.0, 2.5, 1.5, 1.0, 0.5 kcal/mol/Å², respectively, while for membrane leaflet restraints force constants were 5.0, 5.0, 2.0, 1.0, 0.2, 0 kcal/mol/Å². The timestep was set to 1 fs for the first 3 steps, then to 2 fs. We kept planarity of the membrane by restraining the head part of lipids in +19 and -19 Å position compared to the center of the lipid bilayer with a potential force constant of 5.0, 5.0, 2.0, 1.0, 0.2, 0 kcal/mol/Å². The equilibration procedure was followed by 600 ns of free molecular dynamics simulations for each system. Electrostatic interactions are calculated using particle-mesh Eward (PME). A cut-off of 12 Å was used to treat non-bonded interactions. The Nose-Hoover Langevin piston method and a Langevin dynamics method were employed to control pressure and temperature. All molecular dynamics simulations were performed using the NAMD software^40^. For each system under study, four other 90 ns MD simulation replicas were considered, whose starting points were taken from the 600 ns-long simulations at times 300, 405, 510 and 600 ns, producing 32 additional runs. In total, we accumulated around 7.7 microseconds of simulations.

### Structural analysis of the simulations

Analyses of the structural data along molecular dynamics simulations (Root-Mean-Squared-Deviation, Root-Mean-Squared-Fluctuations, distances, …) were carried out with the MDAnalysis Python package^41,42^.

The contraction of the DH domain, defined as the relative position of the FBD and NBD domains, has been calculated along the MD trajectories by measuring the distance between the conserved Arg residue in the flavin binding motif (Arg270 and Arg509 in csNOX5 and hNOX5, respectively) and the conserved Cys residue in the NADPH binding motif (Cys444 and Cys740 in csNOX5 and hNOX5, respectively). This distance, noted DRC, is depicted in Figure 2a. We have defined two geometric parameters, θ and ϕ, to monitor the relative motion of the DH and TM domains during the MD simulations. θ monitors the lateral displacement of the DH domain with respect to the TM domain. It is calculated as the angle between the DH domain at a given time during the MD simulation and the DH domain of the csNOX5 initial model, taken as a reference (thus corresponding to θ = 0°) after a structural superposition of the TM domain. It is computed with the angle_between_domains function of the psico module of PyMOL (The PyMOL Molecular Graphics System, Version 2.4 Schrödinger, LLC.). ϕ is a hinge angle between the two domains. We used common features of the two NOX5 models as reference points: the iron of the hemes (depicted by spheres in Figure 2b) and the center of mass of a structural motif present in the DH domain of both models (colored in cyan in Figure 2b).

To study the influence of the repartition of the lipids on the energetics of the electron transfer, we have defined an asymmetry index *I_as_*, that captures the different repartition of the negatively charged POPG lipids around the hemes in the two leaflets.

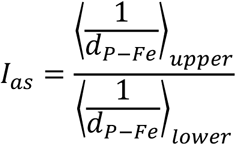

In this equation, *d_P–Fe_* is the 2D-distance in the xy-plane between the POPG molecule’s phosphorus atom and a heme’s iron atom. In the numerator, the average is made over the POPG lipids of the upper leaflet with Heme2, which is located within this upper leaflet (see Figure 1). In the denominator, the average is made over the POPG lipids of the lower leaflet and Heme1. We have used the inverse of the distance as we are interested in the electrostatic interactions between the lipids and the hemes. A value of *I_as_* greater (respectively smaller) than 1 corresponds to a configuration where the POPG lipids in the upper leaflet are on average closer to (respectively further away from) Heme2 than the POPG lipids of the lower leaflet are to (respectively from) Heme1. For a couple of simulations of the same system in two redox states, we compute an average POPG asymmetry index 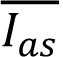 as the average of the instantaneous asymmetry index over all the configurations observed in the two simulations.

### Thermodynamics of inter-heme electron transfer

We have computed inter-heme electron transfer free energies Δ*G* in the framework of Marcus theory. As established by Warshell and co-workers^16^, it is possible to evaluate Δ*G* from two MD simulations performed in the two redox states involved in the electron transfer under study, under the linear response approximation, using:

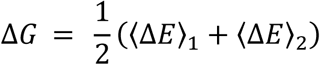

In this equation, Δ*E* = *E*_2_ − *E*_1_ is the vertical energy gap between the two redox states 1 and 2 of energies E1 and E2 respectively. The energy gap is computed and averaged along the simulations in the two redox states to lead to the free energy of electron transfer reaction Δ*G*.

The energy gap can be decomposed in a contribution Δ*E*^*is*^ coming from the redox cofactors (known as inner-sphere contribution), an (outer-sphere) contribution Δ*E*^*os*^ that comes from the interaction of the redox cofactors with the environment, and a coupling term Δ*E^mp^* between them. We rely here on the common assumption that the coupling term is small, so that Δ*G* is the sum of the inner-sphere free energy Δ*G*^*is*^ and the outer-sphere free energy Δ*G*^*os*^. The first term can be computed from quantum mechanics calculations on the isolated cofactors, while the second can be obtained using molecular mechanics calculations. In our case, since the two hemes that are involved in the electron transfer are chemically equivalent, the inner-sphere contribution is zero and the total free energy Δ*G* can be identified with the outer-sphere free energy Δ*G*^*os*^. Since our calculations rely on non-polarizable forcefields, the energy gap can further be decomposed into a sum of contributions from different parts of the system, making it possible to assess their impact on the thermodynamics of the inter-heme electron transfer.

In practice, the Δ*G* values presented here have been computed over the second half of the long molecular dynamics simulations (300 ns) with one evaluation every 100 ps of simulation. For the four 90 ns-long replicas, we used the last 70 ns of simulations to average the energy gaps (again with an interval of 100 ps between 2 energy gap evaluations). Statistical uncertainties have been evaluated using the block average method, considering 5 blocks.

## RESULTS

### csNOX5 and hNOX5 models are structurally stable

We first assessed the structural stability of both models in MD simulations. Figure 3 shows the Root-Mean-Squared-Deviation (RMSD) computed over all atoms of the transmembrane (a) and dehydrogenase (b) domains separately for the eight 600 ns-long molecular dynamics simulations (similar data for the replicas are provided in Figure S2), using the starting point of the simulations as reference. Note that for the DH domain, long and flexible loops (residues 527 to 557, 611 to 632 and 685 to 710 for hNOX5; residues 295 to 301 and 397 to 414 for csNOX5) were removed from the computation of the RMSD. Note also that the longest loop in hNOX5 (residues 527 to 557) is missing in the experimental structure. Our simulation tends to confirm its large conformational variability. In all the simulations the RMSD converges after roughly 300 ns towards values around 2.5 to 3 Å, indicative of a stable structure for both domains. The Root-Mean-Square-Fluctuations of the residues (computed over the backbone atoms), shown in Figure S3 confirm that the secondary structures are stable along the MD simulations and that the conformational dynamics within the TM and DH domains can be explained mainly by the flexibility of loops. This structural stability of the protein is in agreement with what has been observed in previous simulations of csNOX5 and hNOX5^8,12^.

**Figure 3.**
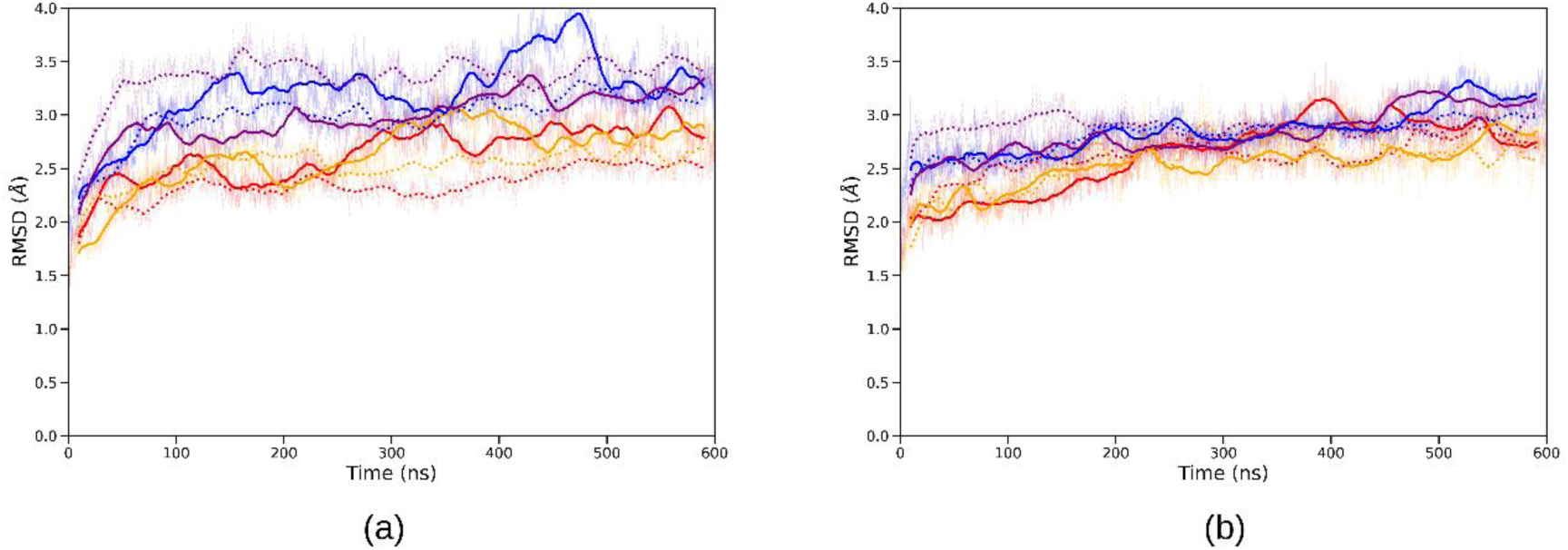
RMSD calculated over the atoms of the transmembrane domain (a) and dehydrogenase domain (b) for the 600ns-long molecular dynamics simulations of systems csNOX5 (red for membrane mbW and orange for the membrane mbH) and hNOX5 systems (blue for membrane mbW and purple for membrane mbH). Solid lines correspond to simulations on the initial redox state (state 1) while dotted lines correspond to simulations in the final redox state (state 2). Transparent lines are for raw data while thick lines are rolling averages over 20 ns.

Figure 4 displays the evolution of θ, that measures the lateral displacement of the DH domain with respect to the TM domain. The initial deviation between the csNOX5 and the hNOX5 models is around 50° (an illustration of this deviation is provided in Figure S4 in Supplementary information). During the simulations, θ is quite stable, with magnitude of variations on the order of 10 to 20 degrees at most. Therefore, no major rotational motion of the DH domain with respect to the TM domain is observed at the microsecond timescale. Variations of the hinge angle ϕ along the MD simulations are shown in Figure S5. The values are about 150° on average and range between 135° and 160° for all the simulations. No correlation of the angle with the model, the membrane composition or the redox state is observed.

**Figure 4.**
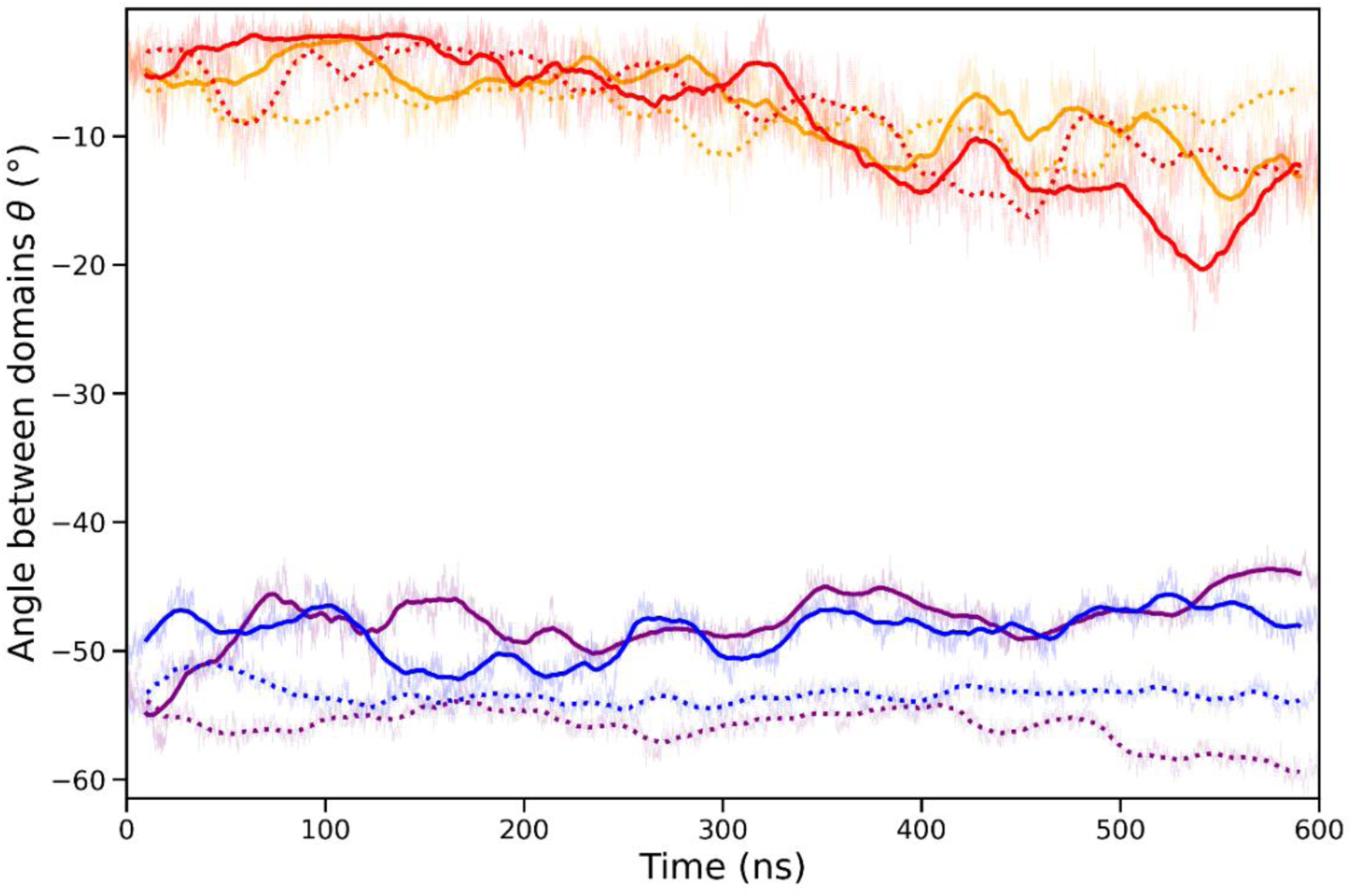
Lateral orientation of the DH domain with respect to the TM domain. The color code and line representations are the same as in Figure 3.

As already stated, the contraction D_*RC*_ of the DH domain is different in our initial models of hNOX5 (18.7 Å) and csNOX5 (13.1 Å). Figure 5 shows the evolution of the contraction of the DH domain along the MD simulations. In hNOX5 simulations, D_*RC*_stays stable with fluctuations on the order of 1 Å, and close to the value found in the inactive state of the NOX2 experimental structure (dark green dashed line). The fluctuations observed in csNOX5 simulations are somewhat larger (around 2-3 Å of magnitude) but overall D_RC_stays close to its initial value and to the one found in the structure of the resting state of NOX2 (light green dashed line).

**Figure 5.**
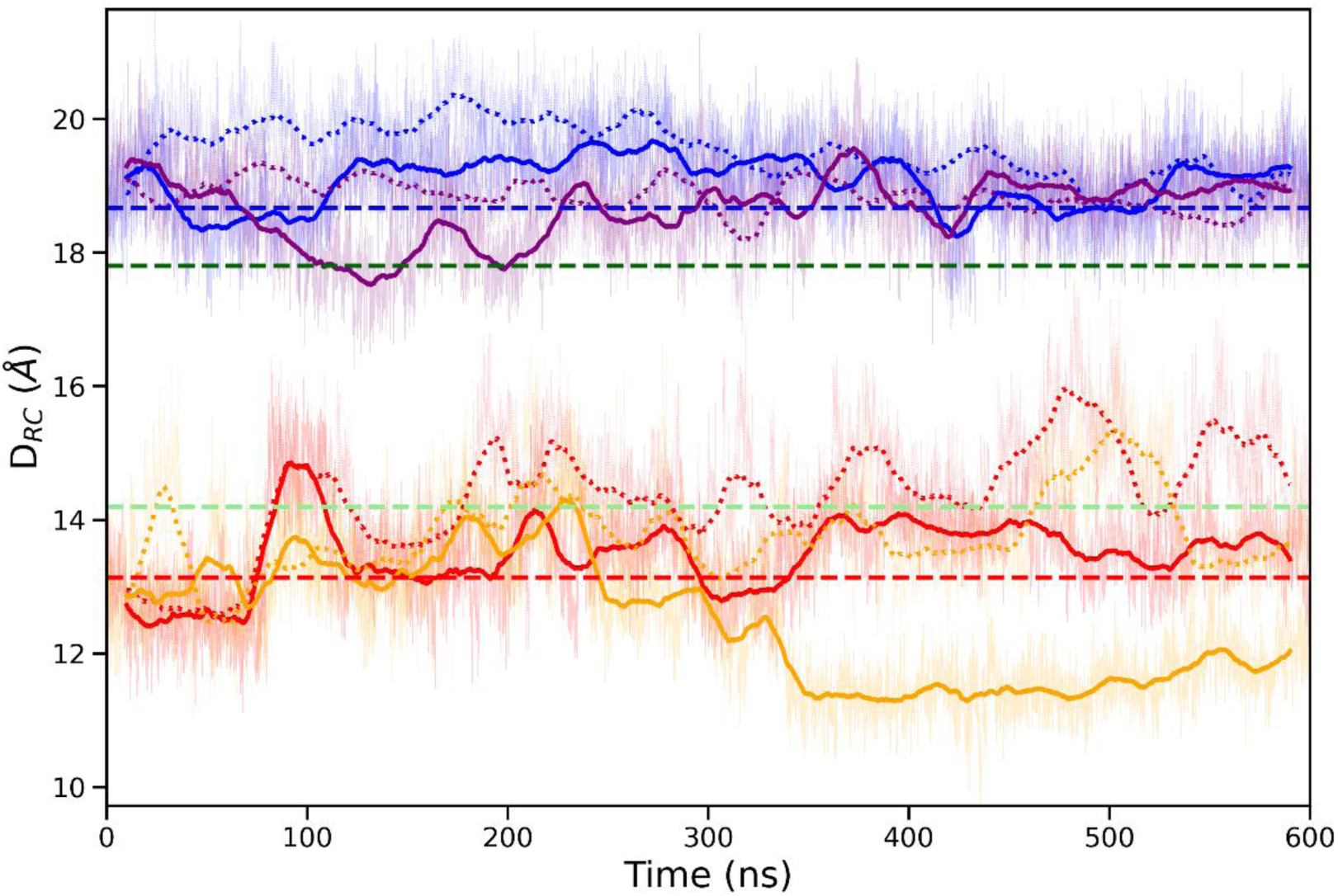
Contraction *D_RC_* of the DH domain along the MD simulations. The color code is the same as in Figure 3. Blue and purple curves correspond to hNOX5 simulations, whereas red and orange curves correspond to csNOX5 simulations. The dashed blue (respectively red) horizontal bar displays the value of D_RC_ in the initial model of hNOX5 (respectively csNOX5). Dashed green bars are the same measures in the NOX2 experimental structures^11^ in the activated (light green) and resting state (dark green)

### Flavin binding differs in hNOX5 and in csNOX5 models

The stability of the binding of flavin has been monitored during the MD simulations. The upper graph of Figure 6 shows the evolution of the RMSD of the isoalloxazine moiety of the flavin cofactor along the MD simulations, the structures having been superposed on the protein atoms. For all but one simulation, this RMSD remains around 3 Å. This value indicates that for almost all the simulations, the cofactor stays close to its initial position in the model. The internal structure of the flavin is however not always maintained and we observed great flexibility of the flavin during the MD simulations, as shown for example in Figure S6 that displays the evolution of the angle made by the adenosine, the ribitol/phosphate chain, and the isoalloxazine moieties of the flavin. The isoalloxazine ring position being quite fixed in the binding site indicates that the adenosine moiety is free to move. We show in Supplementary Information (Figure S7) snapshots taken at the beginning and at the end of the 600 ns-long simulations to illustrate the motion of the flavin.

**Figure 6.**
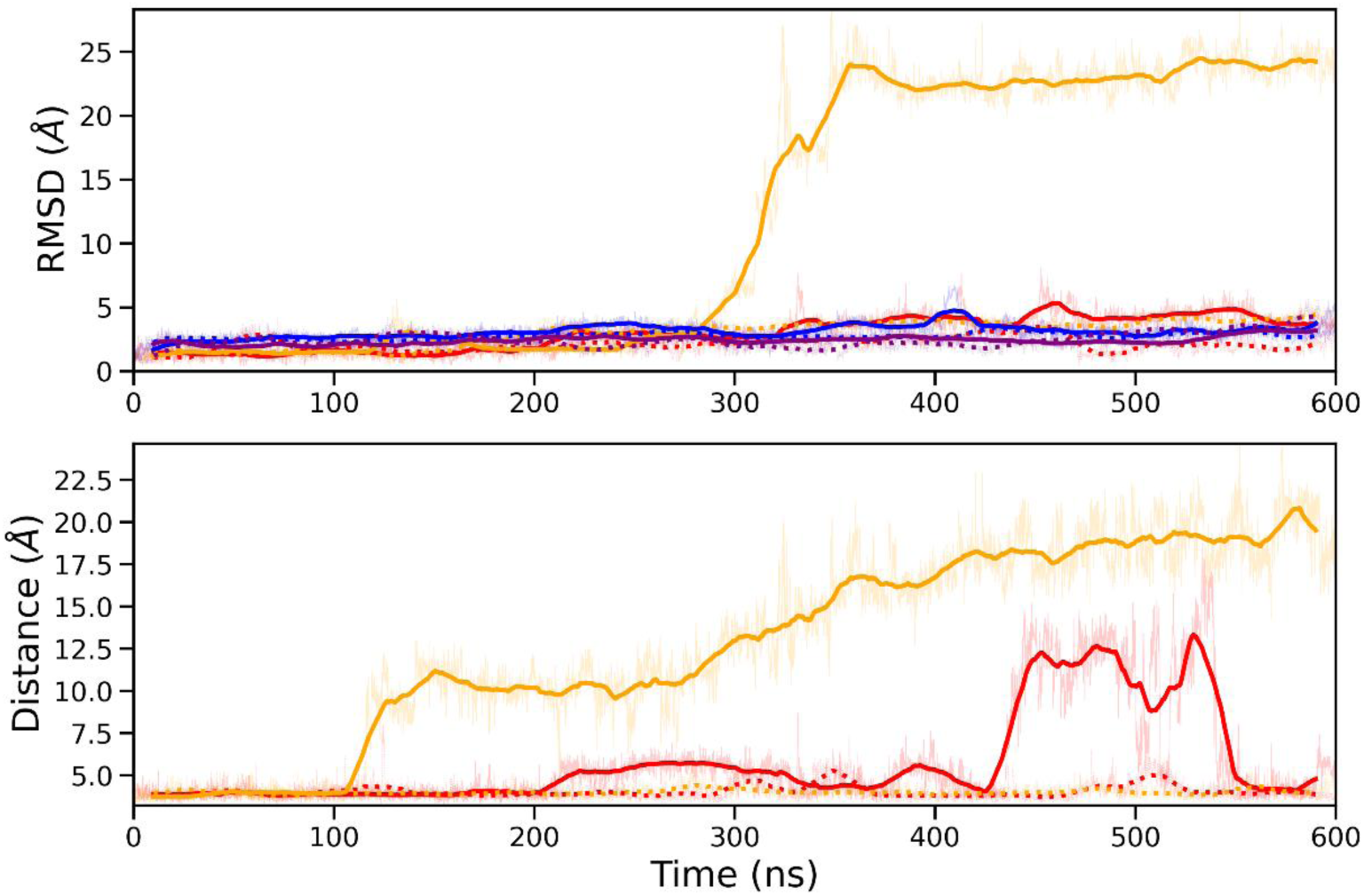
Flavin binding to NOX5 models. The upper panel shows the RMSD of the isoalloxazine moiety of the flavin cofactor after a superposition of the structure on the protein atoms. The lower panel shows the distance between the center of mass of isoalloxazine and the center of mass of the sidechain of residue Phe469 in the csNOX5 simulations. Color code and line representations are the same as in Figure 3.

We have sought to identify the interactions at the origin of the stability of the flavin in its binding site. The orientation of the cofactor being very different in the two NOX5 models under study, these interactions differ between csNOX5 and hNOX5 models. In the case of hNOX5, the isoalloxazine ring makes 4 hydrogen bonds with the protein surrounding, with residues Thr493, His507, Arg509 and Thr594 (see Figure 7a). Note that these residues are highly conserved among the various NOX isoforms. Figure 7b shows that these hydrogen bonds are conserved along the MD simulations (they are present between 85% to 97% of the time). The ribitol/phosphate chain and the adenosine moiety of the flavin make some transient hydrogen bond interactions with the surrounding aminoacids. For instance, the ribitol chain has some interactions with His490 and Pro491, while the adenosine moiety makes some hydrogen bonds with Gly512, Gln513 or even Trp514 and Thr515. Statistics of those interactions are given in Figure S8a.

**Figure 7.**
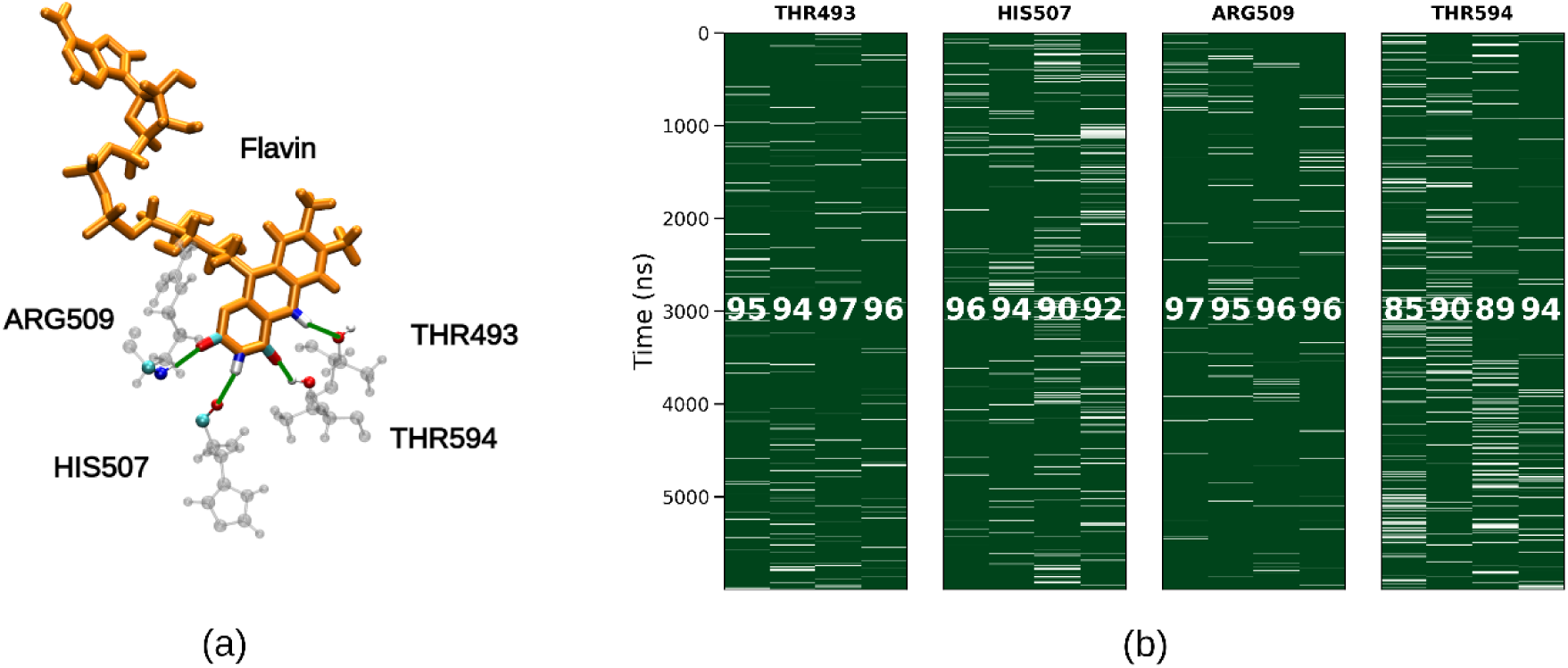
(a) Stable hydrogen bond interactions made by the isoalloxazine ring with the surrounding aminoacids in the hNOX5 structure. (b) Stability of the hydrogen bonds during the MD simulations. For each of the hydrogen bonds, 4 columns are displayed corresponding to the 4 simulations (hNOX5_mbW_st1, hNOX5_mbW_st2, hNOX5_mbH_st1, and hNOX5_mbW_st2). A green color indicates the presence of the hydrogen bond and the number displayed in white in each column is the fraction of simulation time (in percent) during which the hydrogen bond is present.

In the csNOX5 model, the orientation of the flavin is totally different, with the isoalloxazine group facing Heme1. In this orientation, no hydrogen bond is possible between the isoalloxazine group and the protein surrounding. Hydrogen bonding may happen with one propionate from the heme but this interaction is fluctuating over time (with an average of 46% of the time in the csNOX5 simulations and less than 10% of the time in the simulations of hNOX5), and does not provide a decisive contribution to the stability of the binding of the flavin. On the other hand, there is a π-stacking interaction between the isoalloxazine ring and the side chain of Phe469, the C-terminal residue of the DH domain. This interaction is a typical interaction stabilizing flavins in flavoproteins and present in other experimental structures of NOX, such as NOX2 and SpNOX, but not in the hNOX5 experimental model (and absent throughout the MD simulations). The lower panel of Figure 6 shows the distance between the centers of mass of the isoalloxazine ring and of the aromatic ring of Phe469 in the csNOX5 600ns-long MD simulations. In two simulations out of 4, it deviates largely from 4Å, showing the loss of this π-stacking interaction. In the hNOX5_mbW_st1 simulation (red solid line), this loss of interaction happens between roughly 450 and 550ns. In this interval of time, the phosphate groups of flavin find stabilizing interactions with residues Arg270 and Cys444, leaving time for the π-stacking interaction to reform. In the hNOX5_mbH_st1 simulation (orange solid line), the π-stacking is definitely lost after around 150ns of simulations, leading to the detachment of the flavin from its binding site at around 300ns (Figure 6 upper panel and Figure S7b). Once the flavin leaves the binding pocket, the Arg270 and Cys444 sidechains form a hydrogen bond, leading to a contraction of the DH domain (Figure 6). In csNOX5 simulations, there are even less hydrogen bonds formed by the ribitol/phosphate chain or the adenosine moiety than in the hNOX5 simulations (see Figure S8b), adding further lability of the flavin.

### Inter-heme electron transfer is slightly favorable and differs between hNOX5 and csNOX5

Inter-heme electron transfer energetics have been studied in the framework of the Marcus theory. Energy gaps are given in Figure S9 and the resulting free energies are presented in Figure 8. For the four systems based on the two species and two membrane compositions under study, we present 6 values of Δ*G* : one for each of the 90 ns-long replicas (4 first bars), their average, and one value from the second half of the 600 ns-long MD simulations (last bar). The different values are consistent for a given setup. Considering the uncertainties, the inter-heme ET is found to be slightly thermodynamically favorable, with values ranging roughly from -0.25 to -0.05 eV.

**Figure 8.**
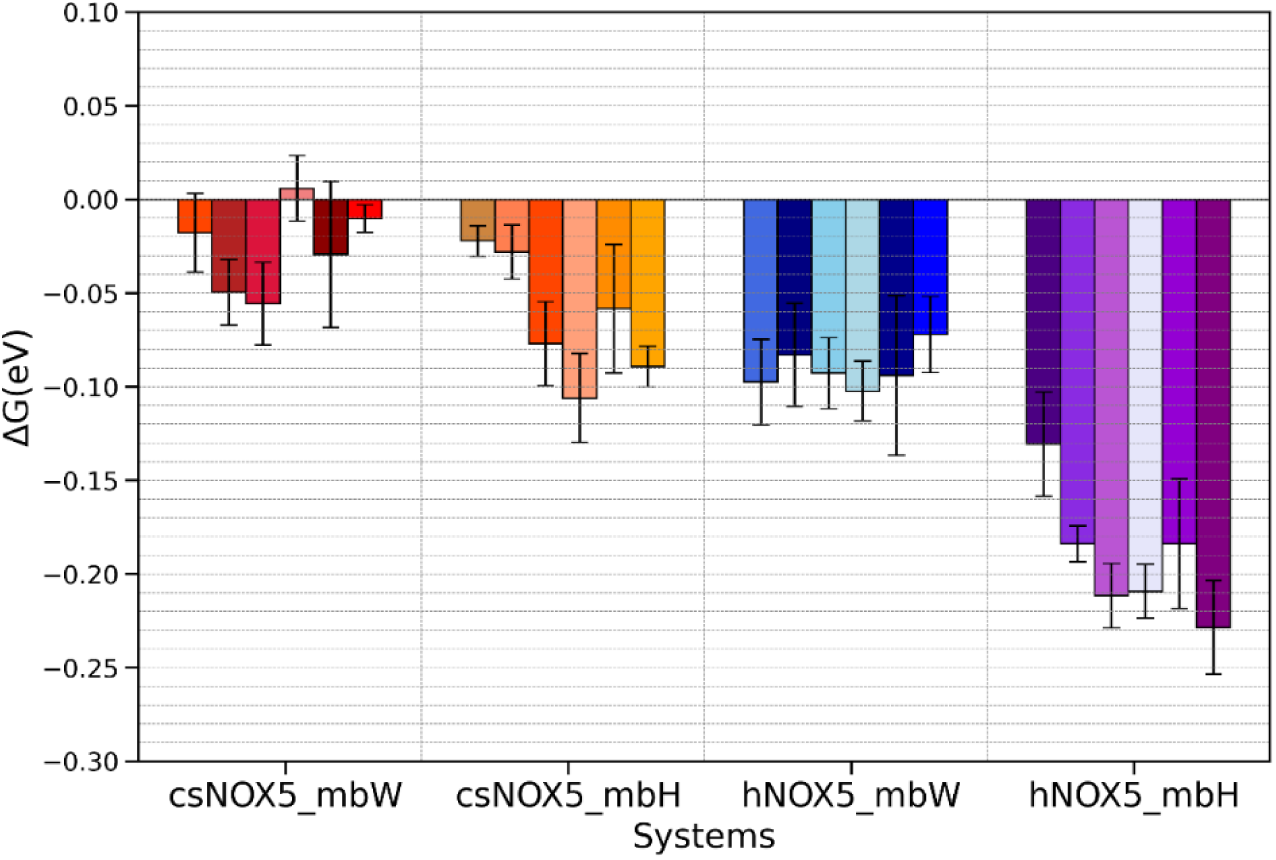
Free energy of inter-heme electron transfer. For each system, six values are presented: the four first values are calculated on the replicas, the fifth value is the average over the replicas, and the last value is computed on the 600 ns-long MD simulations. Error bars correspond to one standard deviation. More details are given in the Computational Methods section.

Two global conclusions can be drawn from the values in Figure 8 when comparing the 4 setups: (i) the increase of the concentration of anionic lipids in the membrane leads to a slight increase, around 0.05 and 0.1eV for csNOX5 and hNOX5, respectively, of the absolute Δ*G* values and (ii) inter-heme ET is more favorable in hNOX5 than in csNOX5 with a difference of roughly 0.1 eV.

A first decomposition of Δ*G* in various contributions is given in Figure 8. The flavin cofactor has a negative contribution of around –0.7 eV in all systems. This trend is probably due to the negative charge borne by the phosphate groups, favoring the electron to lie on Heme 2 located further away. The localization of the neutral isoalloxazine ring of FADH, much different in the hNOX5 and csNOX5 models, plays a negligible role, as it does not significantly modulate this value. The protein contribution (TM and DH domains), on the contrary, largely disfavors electron transfer. It is quite similar for the TM domain but differentiates in the DH domain (see discussion below), where its contribution is even more positive in hNOX5 than in csNOX5. This apparent contradiction with the fact that the total free energy is lower in hNOX5 than in csNOX5 is explained by the opposite, partly compensating contribution of the membrane: positive in the case of csNOX5 and negative in hNOX5. We will come back to this point in the following sections. The aqueous environment, comprised of water molecules and counterions, is always largely favorable to the ET, compensating for the protein and sometimes membrane, contributions.

The contribution of the protein can be further decomposed into the contribution of the transmembrane and the dehydrogenase domains. The former is similar in all the systems, ranging from 1.3 to 1.6 eV. The difference between csNOX5 and hNOX5 thus resides in the DH domain contribution. In csNOX5, it is slightly positive whereas it is much higher (1.5 to 2 eV) in hNOX5. A simple explanation can be given by comparing the aminoacid sequence of the DH domain in the two species (Figure S1): the total charge of the DH domain is +5 (19 Lys, 11 Arg, 10 Asp and 15 Glu) for csNOX5 whereas it is +20 (28 Lys, 21 Arg, 12 Asp and 17 Glu) for hNOX5. Transfer of the electron from Heme1 to Heme2, thus further away from the DH domain, is electrostatically disfavored because of the positive charge of the DH domain, especially in the case of hNOX5. One can note that in the more anionic membrane mbH, the DH contribution is more positive, whereas the TM domain contribution is not sensitive to the membrane composition.

### POPG repartition in the membrane affects inter-heme electron transfer

We now discuss the contribution of the lipid membrane to the electron transfer free energy. The membrane is composed of two leaflets, each made of neutral POPE and anionic POPG lipid molecules, enabling us to further decompose this term into four contributions (see Figure S11). POPE lipids have a negligible effect on the thermodynamics of the electron transfer, less than 0.2 eV. On the contrary, contribution of the POPG lipids of a single leaflet is large in magnitude, around 2-3 eV for the weakly charged membranes mbW and 8-10 eV for the highly negatively charged membranes mbH, and of opposite sign in the two leaflets. Overall, this distribution leads to only a mild contribution of the membrane, constrained between –1 and +1 eV. This almost complete compensation of the contribution of the two leaflets takes its origin in the position of the two hemes inside the membrane. Heme1 is approximately positioned in the middle of the lower leaflet while Heme2 lies in the middle of the upper leaflet, as can be observed in Figure 1.

Understanding the different effect of the membrane in the four systems under study however requires reaching a finer level of detail. Figure 10a shows the contribution of the negatively charged POPG lipids to the energy gaps Δ*E_POPG_* as a function of the membrane POPG asymmetry index *I_as_*, defined in the Computational Methods section, computed over the configurations of the 600 ns-long MD simulations. Same data, aggregated over all the MD simulations is provided in Supplementary information (Figure S12). For a given system, a clear positive correlation is obtained between the two quantities in each system (coefficient of determination *R*^2^ of 0.72, 0.82, 0.86, 0.72 for csNOX5_mbW, csNOX5_mbH, hNOX5_mbW and hNOX5_mbH respectively). This correlation can be explained in terms of simple electrostatics. Indeed, due to the vertical localization of the two hemes (Heme1 in the lower leaflet and Heme2 in the upper leaflet), redox state 1 (reduced Heme1 and oxidized Heme2) is favored upon redox state 2, if the environment of Heme2 is more negatively charged than the environment of Heme1. Therefore, a configuration with a high value of the asymmetry index *Ias* will correspond to a high value of Δ*E_POPG_*.

The data for csNOX5_mbH, hNOX5_mbW and hNOX5_mbH follow the same trend but the csNOX5_mbW (in red) is clearly shifted upwards. Although the *Ias* values are in between those of hNOX5_mbW and csNOX5_mbH, the values of Δ*E_POPG_* are much higher. This shift is due to a strong dissymmetry of the numbers of POPG lipids in the 2 leaflets: there are only 31 POPG in the lower leaflet, against 38 in the upper, leading to a relative difference of almost 9%, much larger than in the other systems (4, 3, and 3% respectively in the case of csNOX5_mbH, hNOX5_mbW and hNOX5_mbH – see Table 1). The consequence is that the membrane in the csNOX5_mbW system is globally more favorable to redox state 1 and the energetic gaps are higher than in the other systems for comparable *Ias* values.

Figure 10b summarizes the results obtained on the two redox states and shows the contribution of the membrane to the ET free energy as a function of the average of the asymmetry index 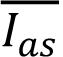. For a given membrane type (mbW or mbH), 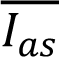 is greater for csNOX5 (red and orange) than for hNOX5 (blue and purple). This increase is due to the largest positive charge of the DH domain of hNOX5 that attracts on average more POPG lipids in the lower leaflet than the DH domain of csNOX5. The increase in the fraction of POPG in the membrane leads to an increase in 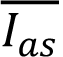 and therefore an increase in Δ*G_memb_*. A linear relation between Δ*G_memb_* and 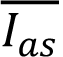 is clearly visible for the systems with the strongly anionic membrane mbH (see the black line on Figure 10) with an almost similar slope of approximately 40 eV for csNOX5 and hNOX5. Such a linear relationship can’t be proven for the weakly charged membranes, because of the large uncertainties on 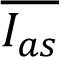 that comes from the low statistics and an expected smaller slope, both due to the smaller number of POPG lipids.

## DISCUSSION

### Inter-heme transmembrane ET thermodynamics in NOX is favorable and finely tuned by the environment

The study of electron transfer in proteins using microscopic simulation emerged in the 80s and is now well established^17,43^. Molecular dynamics simulations are commonly used to investigate the structure and dynamics of membrane proteins or their role to channel water or ions through the membrane^44,45^. The number of studies of electron transfer in membrane proteins inserted in a lipid bilayer has long been limited because of the complexity of the systems involved, although many fundamental processes relying on ET involve membrane proteins such as in respiratory chains, in photosynthetic reaction centers or in NOXs for example. To the best of our knowledge, we trace back the first molecular simulation work on ET taking explicitly into account the membrane to the paper of Tipmanee and Blumberger on electron transfer in cytochrome *c* oxidase^46^. One conclusion of their paper was indeed the necessity to consider the membrane to compute ET thermodynamics parameters for such systems. Since then, the increase in computational power has permitted to tackle the study of electron transfer in large systems inserted into a realistic membrane environment^8,28–32^. For example, Pirnia et al. compared the kinetics of two possible ETs following the initial charge separation in Photosynthetic Heliobacteria Reaction Center using a combination of MD simulations and electronic structure calculations^28^. Few recent studies have focused on proton-coupled electron transfers (PCET) in very large membrane systems (the Alternative Oxidase enzyme^29^, the III2IV2 mycobacterial supercomplex^30^, and Respiratory Complex I^31^) with microsecond-long MD simulations coupled to QM/MM calculations and highlighted the impact of specific residue conformations and protonation states on charge transfer kinetics in these systems. However, to the best of our knowledge, there has been no attempt to characterize in detail the impact of the membrane and its composition on ET thermodynamics, as well as the interplay between the membrane and the other constituents of the system. It is interesting to note also insightful computational studies on multiheme cytochromes from *Shewanella oneidensis* Manganese reducing (MR-1) that sustain extracellular electron transport. The free energy landscape for electron/hole transport can be subtlety modulated by the electrostatic environment and the redox states of the co-factors^47^.

Our results show that inter-heme transmembrane electron transfer is thermodynamically favorable in csNOX5 and hNOX5, in line with our previous calculations performed on csNOX5. We provide evidence that modification of the membrane composition tunes the ET free energy. Although this tuning might be of biological significance, knowing that the phospholipid membrane composition changes during NOX2 activation^26,27,48^, one must keep in mind that the inter-heme ET is only one step within the full mechanism leading to superoxide production in NOX.

It is interesting to note that this modulation in Δ*G* is relatively small, on the order of 0.1 eV, once integrated over all contributors, despite substantial modifications of the membrane content. We could rationalize this result by decomposing the ET free energy (Figure 9). The increase in the POPG:POPE ratio makes the membrane less favorable - or even unfavorable in the case of csNOX5-to the electron transfer but this obstacle is counterbalanced by the aqueous environment that becomes even more favorable, on the same order of magnitude. This compensatory effect is seen when one compares csNOX5 and hNOX5 models. Indeed, the total charge of the DH domain strongly differs between the two species, resulting in a very different contribution of the DH domain to the inter-heme ET free energy: close to 0 in csNOX5 and 1.5-2 eV in hNOX5 (Figure 9). But this distribution in turn leads to a different organization of the POPG lipids in the two leaflets of the lipidic membrane, and in turn to the water and ionic environments, which goes against this first-order effect. The total free energy of the inter-heme ET variation is finally reduced to a few tenths of eV. This compensation mechanism is reminiscent of the standard Le Chatelier’s principle in thermodynamics.

**Figure 9.**
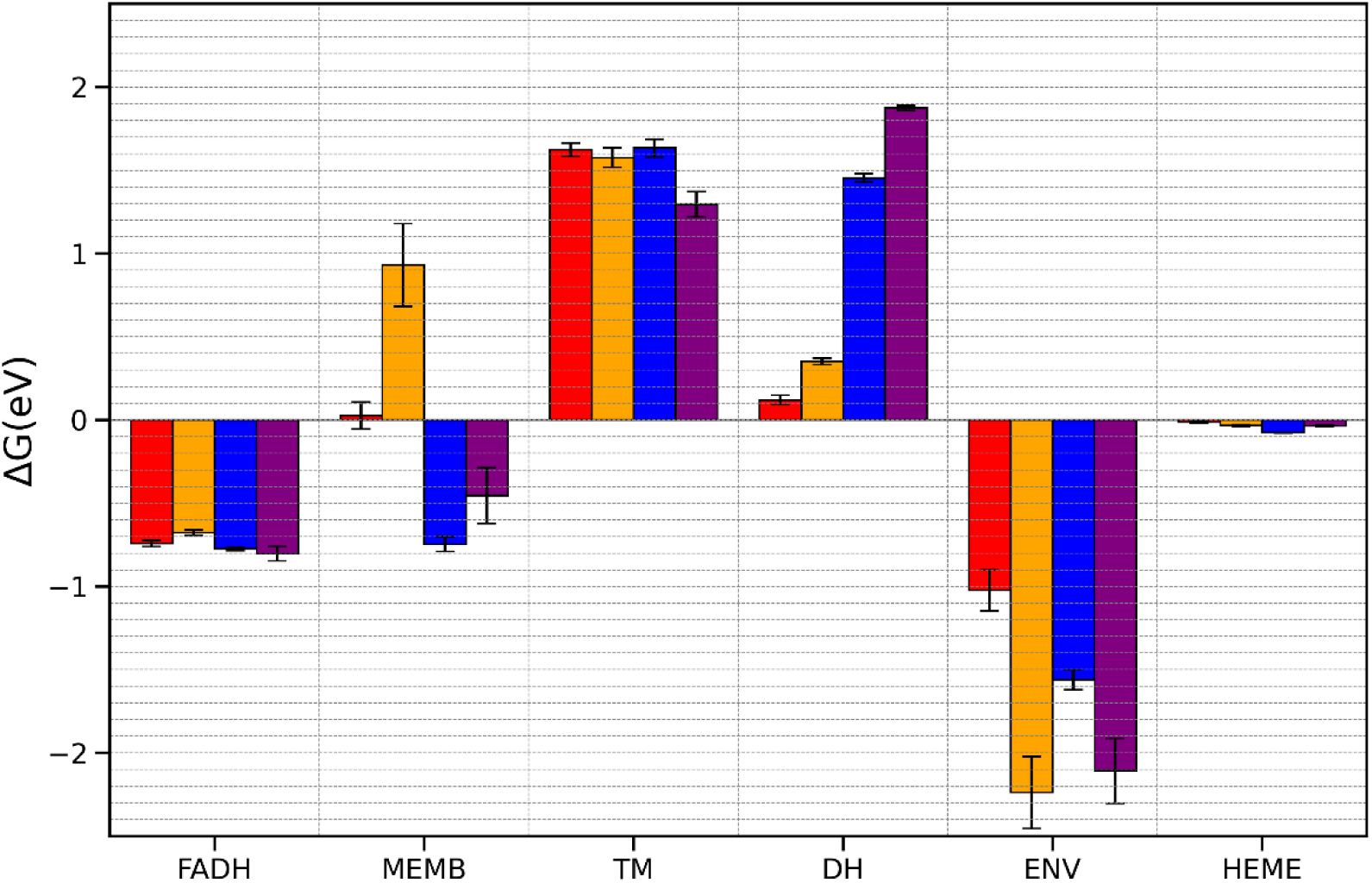
Decomposition of the inter-heme ET free energy into contributions of the different parts of the system: flavin cofactor (FADH), lipidic membrane (MEMB), transmembrane and dehydrogenase protein domains (TM and DH), water and counterions (ENV), and hemes (HEME). The color code is the following: red (csNOX5_mbW), orange (csNOX5_mbH), blue (hNOX5_mbW), and purple (hNOX5_mbH). For clarity, we only present the results for the 600 ns-long MD simulations. Data for the replicas is presented in Figure S10.

**Figure 10.**
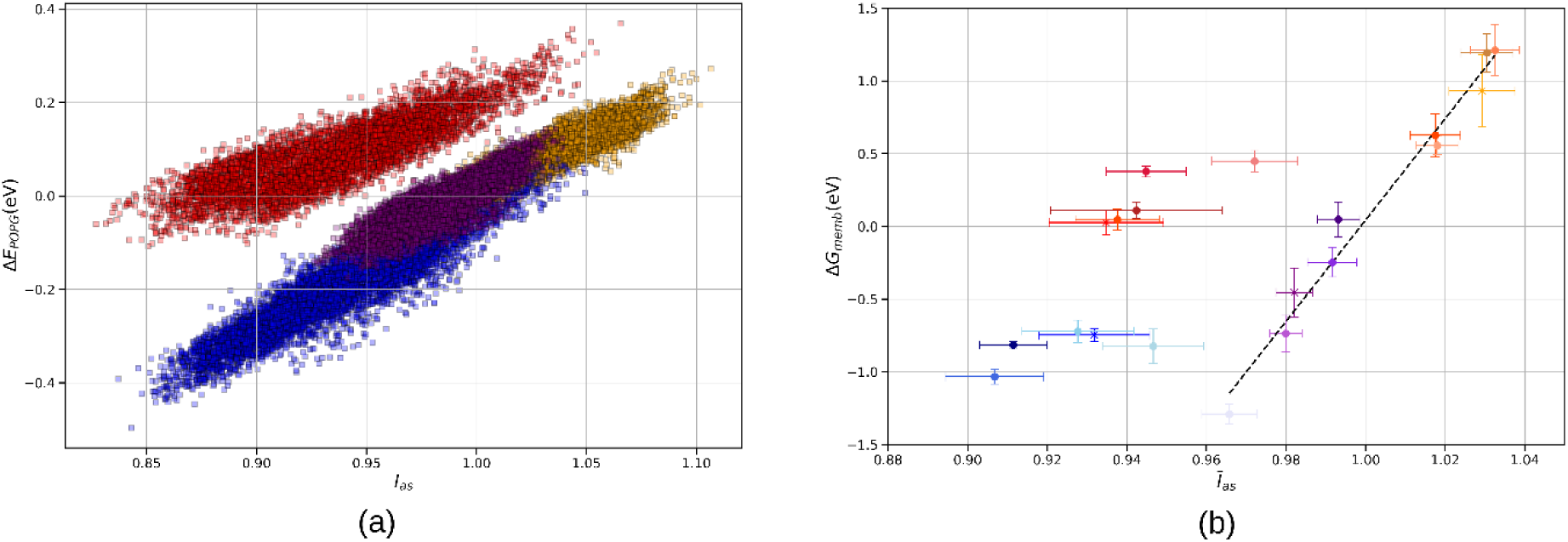
Correlation between lipid repartition and energetics of the inter-heme electron transfer. (a) Contribution Δ*E_POPG_* of the POPG lipids to the energy gap between the two redox states as a function of the membrane POPG asymmetry index Ias. The data shown here has been computed over the second half of the 600 ns-long MD simulations for csNOX5_mbW (red), csNOX5_mbH (orange), hNOX5_mbW (blue) and hNOX5_mbH (purple). Calculations on redox state 1 (respectively state 2) are shown with filled (respectively empty) symbols (b) ET free energy contribution of the membrane Δ*G_memb_* as a function of the average POPG membrane asymmetry index 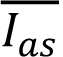. For each system, 5 points are displayed corresponding to the 600 ns-long MD simulations (stars) and the 4 replicas (squares) with color gradient. Horizontal and vertical error bars are computed using block averages with 5 blocks and correspond to one standard deviation.

### Flavin is not always stable in csNOX5 model simulations

In this study, we have extended our previous work based on a model of csNOX5^8^, built as a docked structure between the separated experimental structures of the TM and DH domains^7^. This model exhibits noticeable structural differences with the recent structure of hNOX5^12^ in terms of the conformation of the DH domain, the relative orientation of the TM and DH domains and more importantly the binding of the flavin. The accumulated MD simulation data presented in this paper allows us to better evaluate the biological relevance of this model. The global structure of the protein is stable but flavin binding to csNOX5 appears to be less stable in two simulations (see Figure 6). In this model, the isoalloxazine ring is not maintained by strong hydrogen bonds with the protein, that are found to be extremely stable in the hNOX5 simulations and that are conserved in the experimental structures of other NOX systems. This observation leads us to think that the docked csNOX5 model built in our previous study might not represent a biologically relevant conformation. It remains, however, an interesting model in the context of the study of inter-heme electron transfer in NOX systems, since the contribution of the flavin is rather similar in csNOX5 and hNOX5.

### Is π-stacking with the C-terminal aromatic residue essential for flavin binding?

The presence of an aromatic residue is often observed in the C-terminus of flavoproteins using NAPDH as a substrate such as ferredoxin NADP+ oxidoreductases (FNR), Nitric Oxide Synthase, cytochrome P450 reductase^49^. In many structures, this aromatic residue is stacked to the isoalloxazine ring of the flavin, stabilizing its binding, but at the same time preventing the formation of an NADPH/flavin interaction adequate for hydride transfer. In most NOX proteins, the C-terminal residue is a phenylalanine (Phe765 in hNOX5, Phe469 in csNOX5, Phe570 in hNOX2, Phe397 in SpNOX). Recent models of NOX2 in complex with some cytosolic partners based on alphafold predictions tend to show that this interaction between Phe570 and the isoalloxazine ring may depend on interprotein interactions^50^. Mutation of phenylalanine to serine (which is the C-terminal aminoacid in the constitutively active NOX4 isoform) was shown to significantly decrease flavin binding in SpNOX, but at the same time enhances the catalytic activity^3^. Similar results were obtained in NOX5, where the same mutation results in an almost 3-fold increase in the activity^7^. As suggested by Kean *et al.*^49^, this increase in activity may be explained by a better accessibility of the isoalloxazine ring to the nicotinamide moiety of the NADPH. In NOX2, mutating Phe570 to a glycine or alanine or deleting it leads to a reduced activity^11,51^. Surprisingly, in the hNOX5 experimental structures^12^, the C-terminal phenylalanine residue (Phe765) of the DH domain does not interact with the isoalloxazine ring of the flavin. This lack of π-stacking interaction is consistently observed in all the simulations performed in our work. It would require a complete reorientation of the Phe765 sidechain to reform the interaction, the timescale of which is beyond the length of our simulations. The absence of this interaction does however not affect the stability of the binding of flavin, which is firmly positioned by a network of hydrogen bonds between the isoalloxazine ring and the surrounding aminoacids of the DH domain. This observation may indicate that, as suggested by Liu et al., the role of the C-terminal aromatic residue in NOX may not be to stabilize flavin binding but to help in the formation of a productive conformation, in which the isoalloxazine ring of flavin and the nicotinamide of NADPH come close to each other^11^.

## CONCLUSION

We have reported extensive molecular dynamics simulations in a fully hydrated membrane environment of models of NOX5 from two different organisms (csNOX5 and hNOX5). These models have been inserted in two types of membrane, whose lipid compositions are markedly different in terms of electric charge density to evaluate the impact of the lipidic environment on the inter-heme electron transfer. The csNOX5 model had been built in a previous study and presents structural characteristics that are much different from newer experimental structures, such as the one of hNOX5. Our new data reveals that the binding of the flavin is much less stable in the csNOX5 model than in the hNOX5 one, raising doubt about the biological significance of the csNOX5 binding pose. On the contrary, MD simulations of the hNOX5 model lead to a very stable flavin binding on the timescale of 600 ns.

This work focuses on the study of the inter-heme electron transfer. Our results confirm our previous evaluation of a slightly favorable ET in csNOX5^8^, with free energy of electron transfer ranging from –0.25 to 0 eV. The composition of the membrane thus tunes the thermodynamics of transmembrane ET. Its impact on the other electron transfer steps in NOX has to be evaluated before drawing a conclusion on a possible regulation of ET in NOX by the membrane composition. From a more fundamental point of view, our results highlight the unexpected and previously undocumented multiple dynamical compensatory effects that are at play in complex biomolecular systems.

## ASSOCIATED CONTENT

Supplementary figures and descriptions are given as Supporting Information

## AUTHOR INFORMATION

### Author contributions

FC, AL and MB designed research. BE performed research. All authors discussed and analyzed research data. All authors wrote the manuscript. All authors have given approval to the final version of the manuscript.

## ACKNOWLEDGMENT

We are grateful to Prof. Ji Sun who provided us with the atomic coordinates of the human NOX5 3D model obtained with Cryo-EM experiments prior to publication. We thank Hubert Santuz from Laboratoire de Biochimie Théorique of the IBPC for his help in the use of ColabFold. We thank Dr. L. Baciou and T. Bizouarn for stimulating discussions in the elaboration of this research.

