## Supplementary information for "NOX transmembrane electron transfer is governed by a subtly balanced, self-adjusting charge distribution"

*Baptiste Etcheverry[1], Marc Baaden[2]*, Aurélien de la Lande[1]*, Fabien Cailliez[1]**

1: Institut de Chimie Physique, Université Paris Saclay, CNRS (UMR 8000), 15 avenue Jean Perrin 91405, Orsay, France.

2: Université Paris Cité, CNRS, Laboratoire de Biochimie Théorique, 13 rue Pierre et Marie Curie, 75005, Paris, France

### Sequence alignment of csNOX5 and hNOX5 models

| | |
|---|---|
| csNOX5 | AYIKYYIENNWVKIAFLALYVFVNMFFFMSAVEKYESQGANLYVQIARGCGATLNLNGAL |
| hNOX5 | QLTRAYWHNHRSQLFCLATYAGLHVLLFGLAAS--AHRDLGASVMVAKGCGQCLNFDCSF |
|  | ` : * .*: :: ** *. :::::* *.. :. . * :*:*** **:: ::` |
| csNOX5 | ILIPMLRHFMTWLRKTTINNYIPIDESIEFHKLVGQVMFALAIVHTGAHFLNYTTL---- |
| hNOX5 | IAVLMLRRCLTWLRATWLAQVLPLDQNIQFHQLMGYVVVGLSLVHTVAHTVNFVLQAQAE |
| | `* : ***: :**** * : : :*:*:.*:**:*:* *:..*::*** ** :*:.` |
| csNOX5 | --PIPFAQSLFGTK---------AGISGFLLLLVFIIMWVTAQAPIRKGGKFALFYIAHM |
| hNOX5 | ASPFQFWELLLTTRPGIGWVHGSASPTGVALLLLLLLMFICSSSCIRRSGHFEVFYWTHL |
|  | ` *: * : *: *: *. :*. ***::::*:: :.: **:.*:* :** :*:` |
| csNOX5 | GYVLWFALALIHGPVFWQWVLLPVVGFIIELVIRWKTTKE-PTFVVNASLLPSKVLGLQV |
| hNOX5 | SYLLVWLLLIFHGPNFWKWLLVPGILFFLEKAIGLAVSRMAAVCIMEVNLLPSKVTHLLI |
|  | `.*:* : * ::*** **:*:*:* : *::* .* .:: . :::..****** * :` |
| csNOX5 | QRPQSFNYQPGDYLFIKCPGISKFEWHPF**T**ISSAPEMPDVLTL**H**I**R**AVGSWTGKLYQLIR |
| hNOX5 | KRPPFFHYRPGDYLYLNIPTIARYEWHPF**T**ISSAPEQKDTIWL**H**I**R**SQGQWTNRLYESFK |
|  | `:** *:*:*****::: * *:::************ *.: ****: *.**.:**: ::` |
| csNOX5 | EQREEWI-----------------RSG-----SSQSLPGVPVYIDGPYGTPSTHIFESKY |
| hNOX5 | ASDPLGRGSKRLSRSVTMRKSQRSSKGSEILLEKHKFCNIKCYIDGPYGTPTRRIFASEH |
|  | ` . .* ..:.: .: *********: :** *::` |
| csNOX5 | AILICAGIGV**T**PFASILKSILHRNQQNP---------------AKMPLKKVHFYWLNREQ |
| hNOX5 | AVLIGAGIGI**T**PFASILQSIMYRHQKRKHTCPSCQHSWIEGVQDNMKLHKVDFIWINRDQ |
|  | `*:** ****:*******:**::*:*:. :* *:**.* *:**:*` |
| csNOX5 | KAFEWFVELLSKIEAEDT-----NNLFDLNLYLTGAQQKSDM---------------KV |
| hNOX5 | RSFEWFVSLLTKLEMDQAEEAQYGRFLELHMYMTSALGKNDMKAIGLQMALDLLANKEKK |
|  | `::*****.**:*:* ::: ..:::*::*:*.* *.** * ` |
| csNOX5 | DLITGLKSRTKTGRPDWEEIFKDVAKQHAPDNVEVFF**C**GPTGLALQLRHLCTKYGFGYRK |
| hNOX5 | DSITGLQTRTQPGRPDWSKVFQKVAAEK-KGKVQVFF**C**GSPALAKVLKGHCEKFGFRFFQ |
|  | `* ****::**: *****.::*:.** :: .:*:***** .** *: * *:** : :` |
| csNOX5 | EN**F** |
| hNOX5 | EN**F** |
| | `***` |

***Figure S1: Sequence alignment of the csNOX5 and hNOX5 transmembrane and dehydrogenase domains. Some regions described in the text have been highlighted. Aminoacids highlighted in magenta form big mobile loops observed in RMSF calculation (Supplementary Figure S3) and are excluded from the RMSD calculation on the DH domain (Figure 3 in the main text). Red, Blue, Brown and Green highlighted parts are well conserved motifs between csNOX5, hNOX5 and other NOXs sequences. Residues in bold and underlined correspond to the main residues involved in hydrogen bonds with the isoalloxazine ring of the flavin cofactor (Figure 7 in the main text), the Cysteine used to measure the contraction of the DH domain, and the C-terminal Phenylalanine residue.***

#### Structural stability of the TM and DH domains

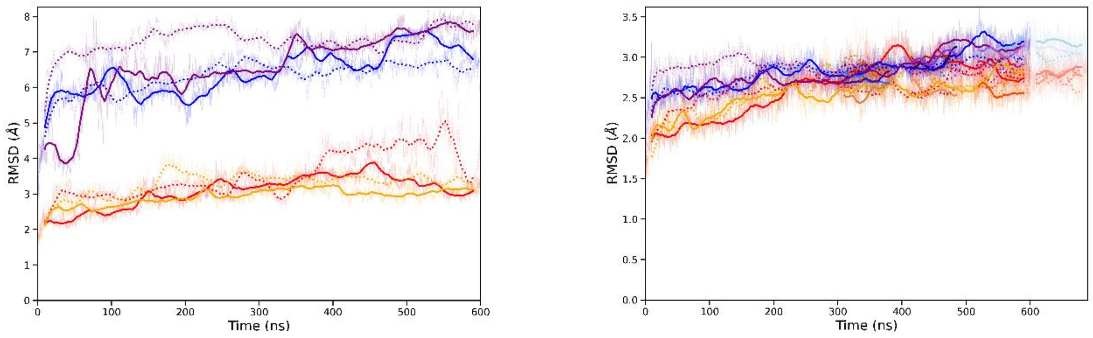

(b)

***Figure S2: (a) RMSD of the DH domain considering the mobile loops in the 600ns simulations. (b) RMSD of the DH domain (without the mobile loops) for all the simulations (the 600ns-long trajectories and the 32 replicas).***

#### Flexibility of loops in NOX5

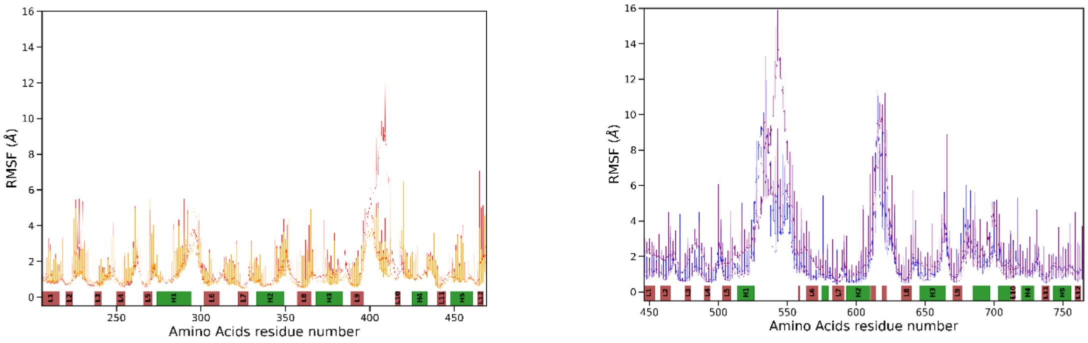

***Figure S3 : Root Mean Square Fluctuations (RMSF) of dehydrogenase backbone atoms in csNOX5 (left panel) and hNOX5 (right panel) 600ns-long MD simulations. The color-code of the lines is the same as in the main text. hNOX5 amino-acid numbering has a shift of about 200 compared to csNOX5, explained by the lack of EF-hand domain in the experimental model of csNOX5. Secondary structures are described at the bottom of the graph by color bars (helices in green and beta sheets in brown). Structural motifs that are present in both models have been numbered accordingly.***

As expected, regions of high mobility (corresponding to high values of the RMSF) correspond to loops. In particular, in the *cs*NOX5 model (left panel), the loop composed of residues 395 to 415 is highly mobile. In the *h*NOX5 model, the longest loops from residue 540 to 560 as well as from residue 610 to 625 are highly flexible. As a consequence, they have been removed in the computation of the RMSD of the DH domain to assess its structural stability (see main text).

#### Relative displacement of DH and TM domains

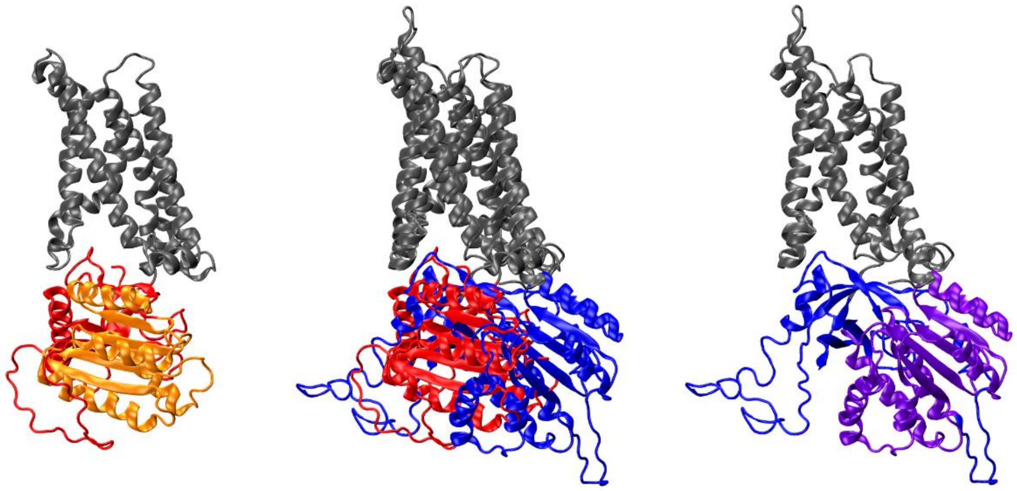

***Figure S4:** Illustration of the different lateral orientation of the DH domain with respect to the TM domain in csNOX5 and hNOX5 models. The two models have been superimposed on the backbone of the first 100 aminoacids of the TM domain. The middle figure shows the superimposed models with their TM domain in grey and the DH domain in red for csNOX5 and blue for hNOX5. The isolated csNOX5 and hNOX5 models are shown respectively on the left and on the right. To better illustrate the position of the domains, the common motif presented in cyan in Figure 2 of the main text is shown in the isolated models in orange for csNOX5 and purple for hNOX5.*

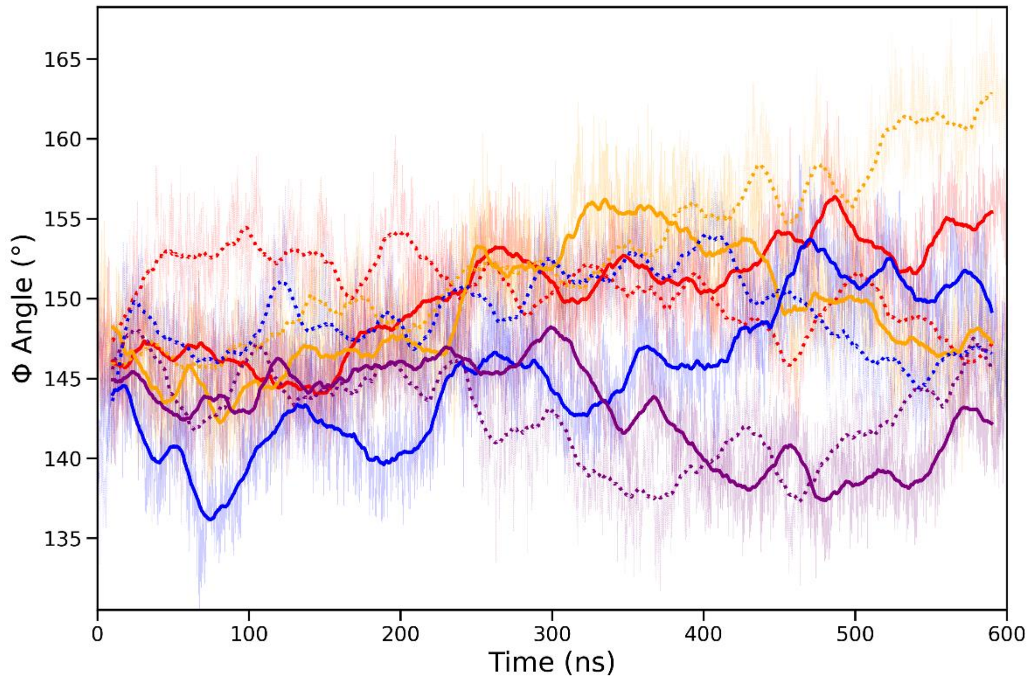

***Figure S5:** Hinge angle $\phi$ between the TM and DH domains. The color-code and line representations are the same as in the main text.*

#### Flexibility of the flavin cofactor structure

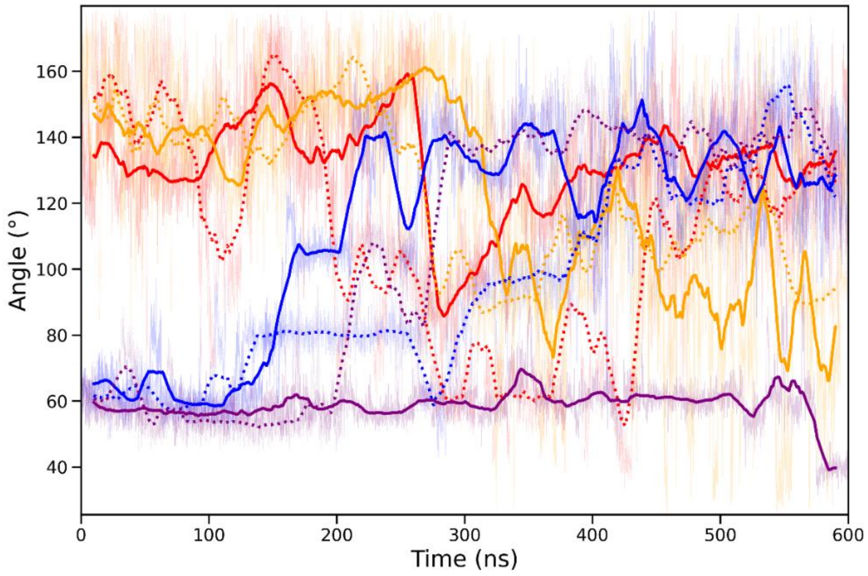

***Figure S6: Flexibility of the internal structure of flavin cofactor. The color code and line representations is the same as in the main text***

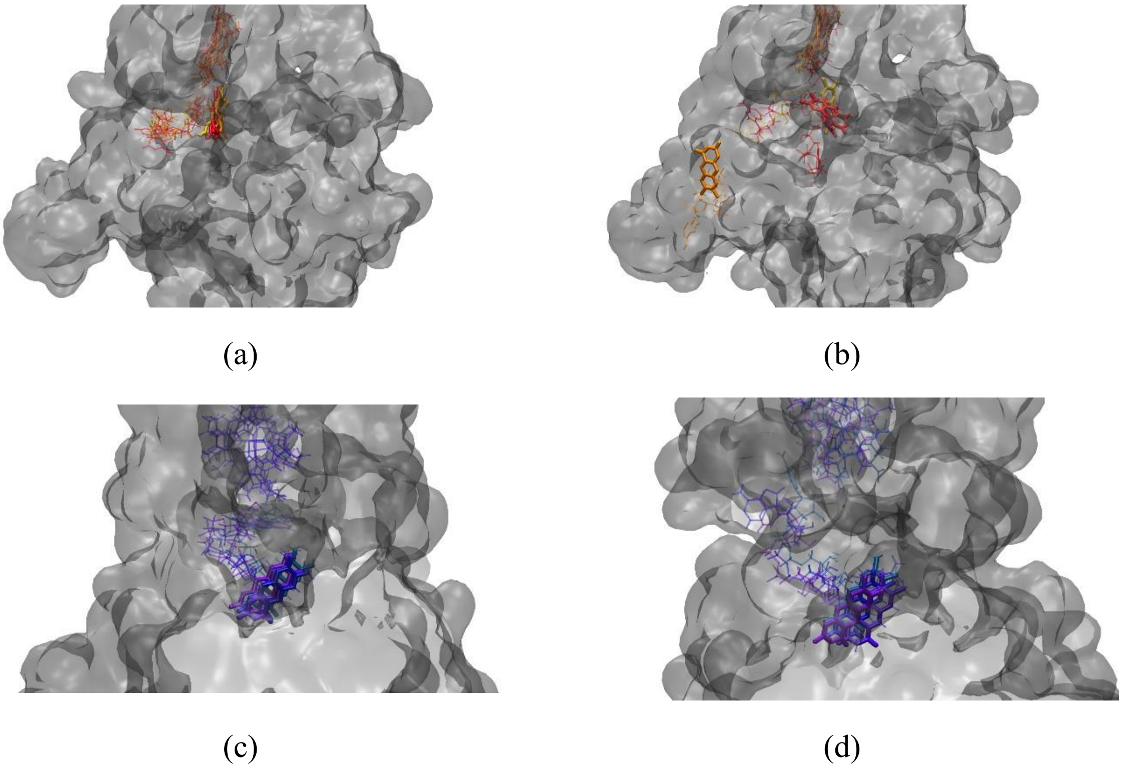

***Figure S7: Representation of flavin binding in the MD simulations at the beginning (a and c) and at the end (after 500ns for b and after 600ns for d) of the 600ns-long MD simulations. The flavin cofactor is shown in licorice representation while the protein surface is represented in grey. (a) and (b) are for csNOX5 simulations with the following color code: red, crimson, orange, yellow for csNOX5_mbW_st1, csNOX5_mbW_st2, csNOX5_mbH_st1 and csNOX5_mbW_st2 simulations respectively. (c) and (d) are for hNOX5 simulations with the following color code: blue, grey-blue, purple, violet for hNOX5_mbW_st1, hNOX5_mbW_st2, hNOX5_mbH_st1 and hNOX5_mbW_st2 simulations respectively***

#### Hydrogen bonds between flavin and aminoacids

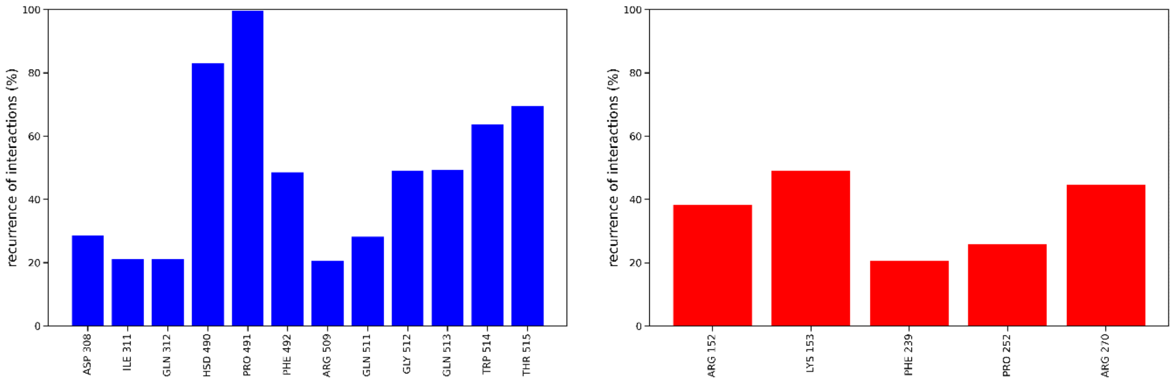

(a) (b)

***Figure S8: Statistics of hydrogen bond interactions between flavin ribitol chain and adenosine moieties and surrounding aminoacids in simulations of hNOX5 (a) and csNOX5 (b). Data is aggregated from the four 600ns-long MD simulations for each system. Only interactions occurring more than 20% of the simulation lengths are shown here.***

#### Energy gaps for inter-heme electron transfer

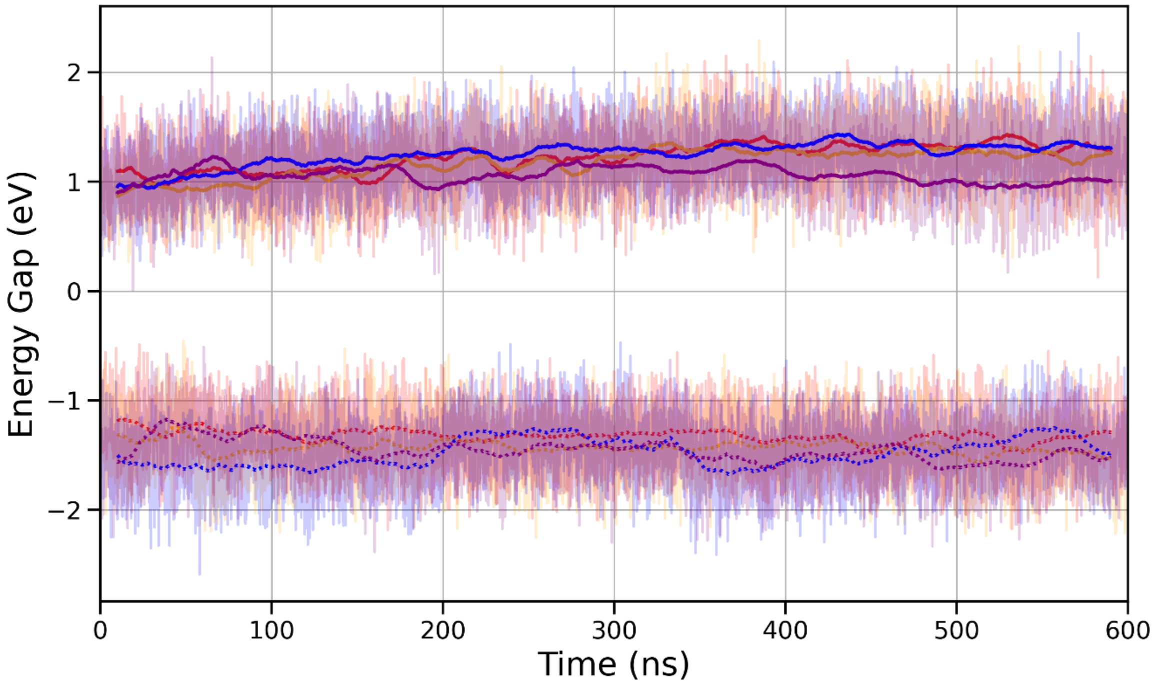

***Figure S9: Energy gap between the 2 redox states computed over the trajectories presented in the main text. The color code and line representations are the same as in the main text. Curves around 1 eV are for simulations on the initial redox state and curves around –1.5 eV are for simulations on the final redox state. Rolling averages over 20ns are shown in thick lines whereas instantaneous energy gaps are shown with transparency.***

#### Decomposition of the contribution of the ET free energy

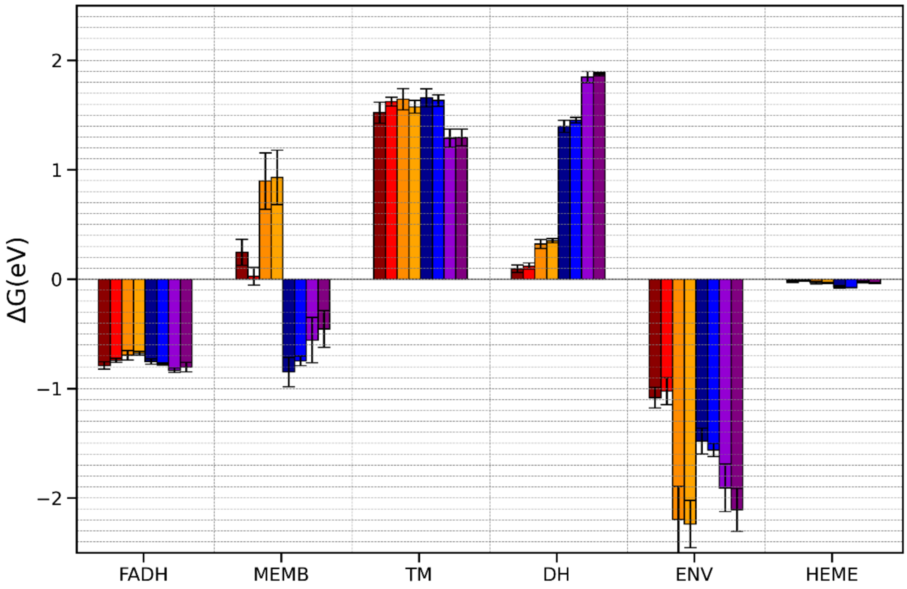

***Figure S10: Decomposition of the inter-heme ET free energy in contributions of the different parts of the system: flavin cofactor (FADH), lipidic membrane (MEMB), transmembrane and dehydrogenase protein domains (TM and DH), water and counterions (ENV), and hemes (HEME). The color code is the following: red (csNOX5_mbW), orange (csNOX5_mbH), blue (hNOX5_mbW), and purple (hNOX5_mbH). Dark colors correspond to the average over the 90ns-long MD replicas and light colors correspond to the 600ns-long MD simulations.***

#### Decomposition of the contribution of the membrane to ET free energy

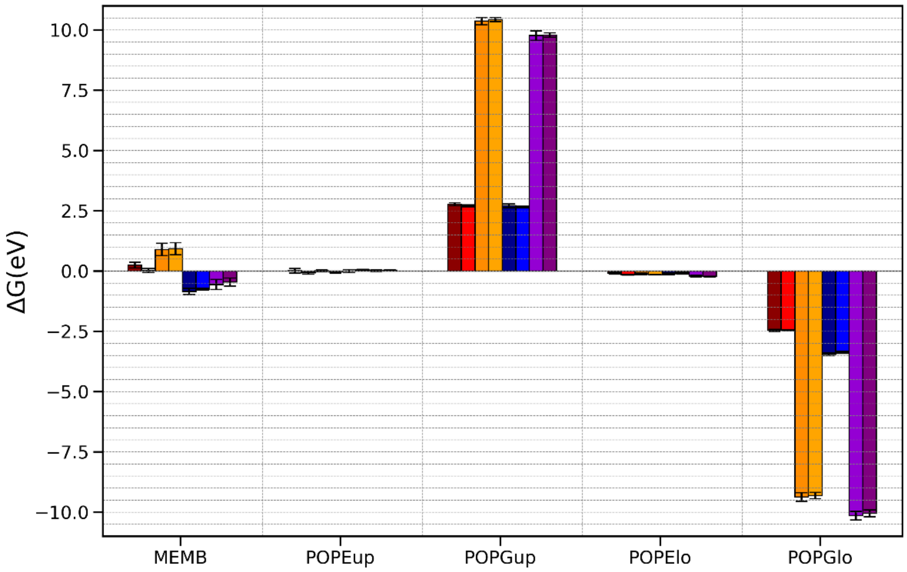

***Figure S11: Decomposition of the contribution of the membrane to the free energy of inter-heme electron transfer. From left to right: full membrane, POPE from the upper leaflet, POPG from the upper leaflet, POPE from the lower leaflet and POPG from the lower leaflet. csNOX5_mbW, csNOX5_mbH, hNOX5_mbW and hNOX5_mbH results are shown in red, orange, blue and purple respectively. Dark color bars (1st, 3rd, 5th and 7th bars of each section) are averages over the replicas. Light bars are the results from the 600 ns-long trajectories.***

### Correlation between POPG lipid repartition and energetics of the inter-heme electron transfer

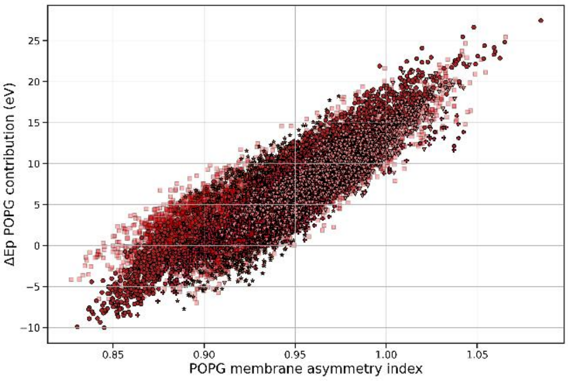

(a)

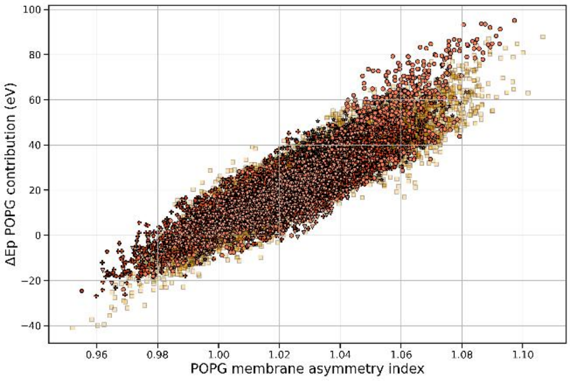

(b)

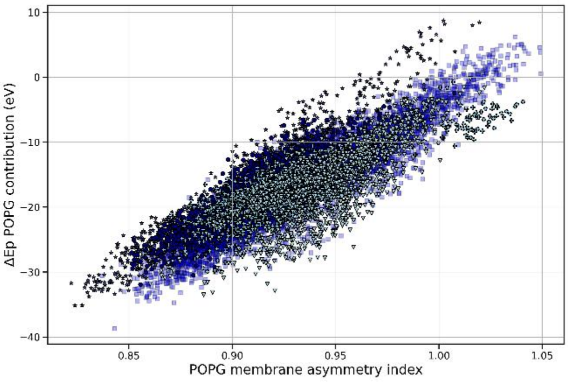

(c)

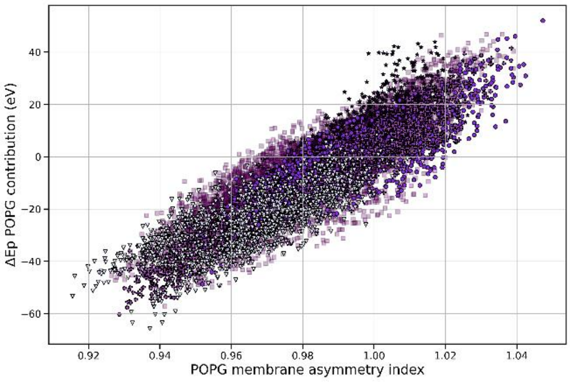

(d)

***Figure S12: Contribution of POPG lipids to the energy gaps as a function of the POPG asymmetry index. Data from all simulations (600ns-long MD trajectories and replicas and in the two redox states) has been aggregated for csNOX5_mbW (a), csNOX5_mbH (b), hNOX5_mbW (c) and hNOX5_mbH (d).***
